# IL-7-adjuvanted vaginal vaccine elicits strong mucosal immune responses in non-human primates

**DOI:** 10.1101/2020.11.09.374405

**Authors:** Sandrine Logerot, Suzanne Figueiredo-Morgado, Bénédicte Charmeteau-de-Muylder, Abdelkader Sandouk, Anne-Sophie Drillet-Dangeard, Morgane Bomsel, Isabelle Bourgault-Villada, Anne Couëdel-Courteille, Rémi Cheynier, Magali Rancez

## Abstract

Mucosal immune responses are crucial in protecting against pathogens entering through mucosal surfaces. However, due to difficulties in disrupting the tolerogenic environment associated with mucosa, mucosal immunity remains difficult to stimulate through vaccines and requires appropriate adjuvants. We previously demonstrated that either administered systemically to healthy macaques or locally expressed in the intestinal mucosa of acutely SIV-infected macaques, interleukin-7 (IL-7) triggers chemokine expression and immune cell homing into mucosae, suggesting its important role in the development of mucosal immune responses.

We therefore examined whether local delivery of recombinant glycosylated simian IL-7 (rs-IL-7gly) to the vaginal mucosa of rhesus macaques could prepare the lower female genital tract (FGT) for subsequent immunization and act as an efficient mucosal adjuvant.

First, we showed that local administration of rs-IL-7gly triggers vaginal overexpression of chemokines and infiltration of mDCs, macrophages, NKs, B- and T-cells in the chorion while MamuLa-DR^+^ APCs accumulated in the epithelium. Subsequent mucosal anti-DT immunization in macaques resulted in a faster, stronger, and more persistent mucosal antibody response compared to DT-immunization alone. Indeed, we detected robust productions of DT-specific IgAs and IgGs in their vaginal secretions and identified cells secreting DT-specific IgAs in their vaginal mucosa and IgGs in draining lymph nodes.

Finally, the expression of chemokines involved in the organization of tertiary lymphoid structures (TLS) was only increased in the vaginal mucosa of IL-7-adjuvanted immunized macaques. Interestingly, TLSs developed around PNAd^+^ high endothelial venules in their lower FGT sampled 2 weeks after the last immunization.

Non-traumatic vaginal administration of rs-IL-7gly prepares the mucosa to respond to subsequent local immunization and allows the development of a strong mucosal immune response in macaques, through the chemokine-dependent recruitment of immune cells, the activation of mDCs and the formation of TLSs. The localization of DT-specific IgA plasma cells in the mucosa argues for their contribution to the production of specific immunoglobulins in the vaginal secretions. Our results highlight the potential of IL-7 as a potent mucosal adjuvant to stimulate the FGT immune system and elicit vaginal antibody responses to local immunization, which is the most promising way to confer protection against many sexually transmitted diseases.

## INTRODUCTION

Mucosae form a physical barrier that limits the invasion of pathogens in the host but also ensures important physiological functions that require a certain degree of porosity. Because of their locations and these two antagonistic characteristics, mucosae are equipped with a peculiar immune system that constitutes a first line of defense for the organism. IgAs have a compartmentalized distribution and repertoire that are believed to contribute to the protection of mucosal surfaces. Strengthening mucosal immunity should be effective in increasing protection against invasive pathogens, but it is difficult to achieve through systemic vaccination.

The administration of a vaccine on mucosal surfaces is a promising way of inducing such immunity, however, it is a method that necessitates adequate adjuvants and is less often explored. The development of such an adjuvant requires understanding the specific mechanisms involved in establishing protective mucosal immunity and adapting the adjuvants to each specific mucosa. Indeed, contrarily to a generally accepted idea, the mucosal immune system certainly does not use common mechanisms to develop immune responses at all sites. Indeed, distinct vaccination routes, oral, nasal, sublingual, rectal or vaginal, stimulate mucosal immunity in different locations (1, 2). Furthermore, vaginal immunization leads to more robust vaginal IgG and IgA antigen-specific antibody responses than parenteral immunization or immunization at other mucosal sites (3, 4).

In the presence of antigens on the mucosal surface, the induction of mucosal immune responses occurs in organized mucosal lymphoid tissues and in draining lymph nodes (LNs). In the mucosa, epithelial cells serve as sensors that detect microbial components through pattern-recognition receptors and transfer signals to underlying mucosal cells to trigger innate, non-specific defenses and promote adaptive immune responses. The signals involved in the differentiation and tissue homing of antigen-specific lymphocytes in the different mucosae remain to be fully defined but, as a whole, this mechanism leads to the preferential development of immune responses at the site where the antigen or the pathogen was initially encountered.

It is known that the tissue-specific expression of chemokines, integrins and homing receptors are involved in immune cells homing to the FGT, however, there is less research conducted on the mechanisms involved in cells homing to the FGT than in the gut or the lungs. Various chemokines were identified to induce cell homing into the vaginal mucosa (CCL2 (MCP-1), CCL5 (RANTES), CCL7 (MCP-3), CCL20 (MIP-3α), CXCL8 (IL-8), CXCL9 (Mig), CXCL10 (IP-10) CXCL12 (SDF-1), CCL28 (MEC)…) (5–12). In addition, α4β7 and α4β1 integrins have been described as participating in the development of vaginal immunity in mice (13, 14). Besides, the expression of the vascular cell adhesion molecule-1 (VCAM-1), which binds to these integrins, has been detected in the human vagina (15) and is overexpressed during inflammatory processes in mice, which suggests a role in cell recruitment into the genital mucosa (16, 17).

Considering the still incompletely described chemokine/integrin network in the genital mucosa, the identification of a strategy to stimulate the physiological expression of this complex network triggered by antigenic stimulation could help the development of an effective mucosal adjuvant. Various cytokines such as GM-CSF, IL-2, IL-12, IL-15 and IL-18 and chemokines such as CXCL8, CCL5, CCL3 (MIP-1α), CCL4 (MIP-1β), CCL19 (MIP-3β), CCL20, CCL21 (6Ckine), CCL25 (TECK), CCL27 (CTACK) or CCL28 have been tested, mainly in mice, as potential adjuvants for the development of mucosal immunity (12, 18–21). So far, these studies have remained largely disappointing. In contrast, thymic stromal lymphopoietin (TSLP) administered nasally together with antigens as well as CXCL9 and CXCL10 intravaginal administration after s.c. immunization acted as a potent mucosal adjuvant in mice (22, 23). Finally, lymphotactin (XCL1) and defensins exerted weak adjuvant activity for mucosal immunity when administered nasally with antigens (24).

Recently we and others have evidenced an overexpression of interleukin-7 (IL-7), a cytokine constitutively expressed by mucosal epithelial cells, in tissues following viral and bacterial infections (25–27). During the acute phase of these infections, this cytokine triggers the mucosal expression of various chemokines, favors integrin and chemokine receptor expression by T-cells and leads to immune cell homing into various mucosal tissues of both humans and rhesus macaques (25, 28, 29). In addition, IL-7 has been shown to play an important role in the formation of tertiary lymphoid organs (30–33). Moreover, IL-7 also contributes to lymphangiogenesis (34). Increased levels of IL-7 observed in infected tissues could thus participate in the induction of antigen-specific immune responses in infected mucosae.

Considering that IL-7, either systemically administered or locally expressed in acutely infected tissues, triggers both the expression of chemokines in tissues and the homing of immune cells into lymphoid and non-lymphoid organs (25, 28, 29), we investigated whether local administration of low doses of IL-7 directly at the surface of the vaginal mucosa could prepare it for subsequent immunization. We evidenced that non-traumatic topical administration of IL-7 triggers major physiological modifications of the vaginal mucosa, characterized by local production of a specific panel of chemokines and infiltration of various immune cells into the chorion. Moreover, we have demonstrated an efficient local immune response in the IL-7-treated vaginal mucosa following local immunization against diphtheria toxoid (DT) used as a model immunogen. Our results emphasize the potential of IL-7, already used in clinics without major adverse effects (35), as a potent mucosal adjuvant to stimulate the FGT mucosal immune system.

## MATERIALS AND METHODS

### Animals, drug administration and tissue collection

The healthy Chinese female rhesus macaques (*Macaca mulatta)* included in this study were housed, cared for, and handled in BSL2 NHP facilities of the Institut Pasteur (Paris, France; accreditation no. A 78-100-3) and IDMIT (“Infectious Disease Models and Innovative Therapies” at the CEA “Commissariat à l’Energie Atomique,” Fontenay-aux-Roses, France; accreditation no. C 92-032-02). Approval number 2010-0008 for the use of monkeys in this protocol was obtained from the ethics committee of Paris 1. All animal handling was carried out under ketamine anesthesia, in accordance with European regulations. The animals were seronegative for SIV_mac_, simian T-cell leukemia virus type 1, simian retrovirus type 1 (type D retrovirus), and herpes virus B.

Recombinant glycosylated simian IL-7 (rs-IL-7gly) was obtained from Cytheris SA (now Revimmune Inc., France) and administered either through intra-mucosal injection at several sites of the vaginal walls (4 injections per macaque), together with black Indian ink (1 to 10 ng/injection site, in 20μL of 1/24 Indian ink, in calcium free Dulbecco’s phosphate buffered saline (PBS), n=8 macaques) or by vaginal spray using the APTAR bidose spray device (1 to 15μg in 200μL of PBS per spray, n=17 macaques). Control animals were untreated or injected with Indian ink alone (n=8 macaques), or sprayed with PBS (n=3 macaques).

Immunization against diphtheria toxoid (DT) was performed through non-traumatic administration of DT (Biological Laboratories, Courtaboeuf, France; 7μg per animal, in 200μL of PBS) into the vaginal lumen, using the APTAR bidose spray device. Immunizations were repeated with an identical protocol at week 16 (boost 1) and week 31 (boost 2) after prime immunization, and all the macaques were euthanized 2 weeks after a third boost immunization performed at week 55 after prime immunization.

Cervico-vaginal lavages (CVL) and blood samples were taken from each animal at baseline and every week throughout the protocol. Each CVL sample was collected using a sterile pipette by inserting in the vaginal cavity 2 mL of sterile PBS which was re-aspirated with the same pipette (12 to 20x), and then added in a sterile 15 mL tube containing antibiotics (200 U/mL penicillin and 200 μg/mL streptomycin, final concentrations) and protease inhibitors used according to the manufacturer’s recommendations (1X of cOmplete ™, EDTA-free Protease Inhibitor Cocktail, Roche Applied Science, Meylan, France). CVLs were centrifuged at 1,800rcf for 1 hour at 4°C then cleared using Spin-X^®^ Tubes centrifuged at 16,000rcf for 30min at 4°C (Sigma-Aldrich, Lyon, France), aliquoted, and stored at −80°C until use.

Vaginal biopsies were taken using biopsy forceps from non-injected healthy animals and at the sites of Indian ink injections (administered alone or together with rs-IL-7gly), 24 or 48 hours after injection, from non-sprayed healthy animals and 48 hours after the administration of rs-IL-7gly by vaginal spray using the APTAR bidose device, as well as 4 weeks before primary anti-DT immunization and 4 weeks after each anti-DT immunization (post-prime and post-boosts n°1 and n°2).

Immediately after sampling, the biopsies were either placed in 600 μL of RLT buffer from the RNeasy kit (Qiagen, Courtaboeuf, France) or snap-frozen in an Optimal Cutting Temperature compound (Tissue-Tek^®^ O.C.T.™ Compound, Labonord, Templemars, France) in isopropanol cooled with liquid nitrogen and stored at −80°C until use.

At necropsy, both the entire vagina and iliac lymph nodes (LNs) were collected and immediately treated for future analyses. Pieces of vaginal tissue (4mm^2^), sampled from the lower and upper parts of the vaginal mucosa or from the vaginal fornix, were either frozen at −80°C in RLT buffer (Qiagen) for future RNA extraction or snap frozen using O.C.T.™ and preserved at −80°C. Pieces of iliac LNs were similarly processed for further analysis. Peripheral blood mononuclear cells (PBMCs) were purified by Ficoll density gradient centrifugation and conserved in liquid nitrogen in fetal calf serum 10%DMSO until use.

### Laser capture microdissection of vaginal mucosal tissue

Twelve-μm thick cryosections of vaginal tissues were collected on a polyethylene foil slide (SL Microtest GmbH, Jena, Germany) and stored at −80°C until use. The cryosections were then air dried for 5 minutes, counterstained with hematoxylin for 30 seconds, air dried, and microdissected as previously described (36). Epithelial tissue and chorion were individually sampled from 3 to 5 consecutive sections and each microdissected tissue was immediately placed in 80μL of ice-chilled RLT buffer (Qiagen) and stored at −20°C until RNA extraction. For each microdissected sample, RNAs were extracted from >2mm^2^ of epithelium and >5mm^2^ of chorion.

### Real-time PCR quantifications

mRNAs were extracted from mucosal biopsies, as previously described (25), or from microdissected samples using the RNeasy tissue kit (Qiagen). Briefly, for microdissected samples, 300 to 400μL of tissue-containing RLT buffer (Qiagen) were extensively vortexed for 3 minutes then centrifuged for 3 minutes (16,000rcf). Residual DNA was removed from the cleared lysate using DNase digestion on columns (Qiagen). mRNAs were eluted in 40μL of RNase-Free water. mRNAs recovered from microdissected samples or vaginal biopsies (about 1 mm^3^) were reverse-transcribed with the QuantiTect Rev Transcription Kit (Qiagen), used according to the manufacturer’s recommendations, and cDNAs were stored at −20°C until use.

The cDNAs were PCR amplified in a final volume of 50μL. PCR amplification consisted of an initial denaturation of 15 minutes at 95°C, followed by 22 (biopsy samples) or 28 (microdissected samples) cycles consisting of 30 seconds at 95°C, 30 seconds at 60°C, and 3 minutes at 72°C using outer 3’/5’ primer pairs.

Multiplex PCR amplifications were optimized to allow simultaneous amplification of (i) CCL3, CCL11, CCL25 and CXCL8, (ii) CCL5, CCL20, CCL28 and CXCL10, (iii) CCL19 and CCL21, (iv) CCL2 and CX_3_CL1, (v) CCL4, (vi) CCL7, (vii) CCL8, (viii) CCL17, (ix) CCL22, (x) CXCL12, (xi) CXCL13, (xii) CD132 and CD127, (xiii) IL-17A and IL-21, (xiv) TSLP, (xv) LTα, and (xvi) LTβ, together with the hypoxanthine phosphoribosyl transferase (HPRT) gene, used as a housekeeping gene. These PCR products were diluted 1/100 in water and used to individually quantify each of the chemokines, CD132, CD127, TSLP, IL-17A, IL-21, LTα, LTβ, or HPRT amplicons, in LightCycler^®^ experiments using inner 3’/5’ primer pairs, as previously described (25). The results were expressed as absolute numbers of target mRNA copies per HPRT mRNA copy. All the primers used in this study are described in **Supplementary Table 1**.

### Immunohistofluorescent staining

Four-μm thick tissue sections fixed with formaldehyde and embedded in paraffin (FFPE) and eight-μm thick cryosections collected on glass slides (SuperFrost® Plus, Menzel-Gläser, Illkirch, France) were immunostained as described in the supplementary material. The antibodies used for immunohistofluorescence labeling are listed in **Supplementary Table 2**.

### Reverse immunohistofluorescent staining

Reverse immunohistofluorescent staining was used to detect cells producing DT-specific antibodies. Ten-μm thick cryosections were fixed for 20 minutes at 4°C in 2% PFA and rinsed with PBS, permeabilized with 0.2% triton for 8 minutes, rinsed with PBS, blocked with 5% BSA for 30 minutes in PBS, then a Streptavidin/Biotin Kit was used according to the manufacturer’s recommendations (Vector Laboratories).

Tissue sections were incubated with rabbit anti-IgA or anti-IgG antibodies (DAKO) for 2 hours at RT, rinsed in PBS/0.5%Tween20, then incubated overnight at 4°C with DT Ag (15μg/mL). Sections were rinsed in PBS/0.5%Tween20, incubated with goat anti-DT-FITC antibodies (Abcam) overnight at 4°C, then rinsed in PBS/0.5%Tween20, incubated with donkey anti-goat Biotin (Abcam) secondary antibodies, rinsed in PBS/0.5%Tween20, incubated with streptavidin-Alexa Fluor^®^ 488 (Molecular Probes) in the dark for 15 minutes at RT, rinsed in PBS/0.5%Tween20, then blocked 30 minutes in 10% normal goat serum and 5% BSA in PBS. IgA or IgG staining were revealed with goat or donkey anti-rabbit-Alexa Fluor^®^ 546 secondary antibodies (Molecular Probes) in the dark for 30 minutes at RT.

The tissue sections were rinsed in PBS/0.5%Tween20, then in PBS alone, counterstained with DAPI (Molecular Probes) and mounted in Fluoromount-G (Southern Biotechnology). The antibodies used for reverse immunohistofluorescence labeling are listed in **Supplementary Table 2**.

As controls for specificity, the anti-DT-FITC antibodies in combination with the Biotin-labeled anti-goat antibodies plus the streptavidin-Alexa-Fluor^®^ 488 did not stain either DT-coated vaginal mucosae or LNs from unimmunized animals, nor uncoated tissues of immunized macaques.

### Image capture and analysis

The immunostained sections were examined under an inverted epifluorescence Leica microscope (DMI6000, Leica Microsystems Gmbh, Wetzlar, Germany), equipped with an ORCA-Flash4.0 LT camera (Hamamatsu Photonics) and coupled with video imaging using the MetaMorph 7.8.8.0 software (Molecular Devices, Sunnyvale, CA, USA). Images were acquired digitally with a 10x or a 20x objective (Leica), then we used both Photoshop (CS5 version, Adobe Systems Incorporated), and ImageJ (1.52p version) software to analyze the stainings. For color images, brightness and contrast were adjusted on each entire digitally acquired image, with the same levels for each labeling set, using the Brightness/Contrast command in Photoshop software.

Quantifications of cells and chemokines in tissue were performed on image reconstructions of the entire sections of the different vaginal biopsies. Sections were digitally acquired at the best focus with a 20x oil objective (Leica using the Yokogawa CSU X1 Spinning Disk (Yokogawa, Tokyo, Japan) coupled with a DMI6000B Leica microscope with MetaMorph 7.7.5 software, using Scan-Slide option (10% overlap). Analyses were performed using an ImageJ routine provided by T. Guilbert (Institut Cochin, Paris, France).

Labeling of immune cells and chemokines was quantified in manually defined zones of chorion or epithelium. After a denoising process, manual threshold was applied on CD3^+^, CD4^+^, CD8^+^, CD20^+^, CD11c^+^, DC-SIGN^+^, PM-2K^+^, MamuLa-DR^+^, CD83^+^ stainings to identify immune cell surfaces, and on CCL2^+^, CCL5^+^, CCL7^+^, CCL19^+^, CXCL12^+^, and CXCL13^+^ stainings to define chemokine expression surfaces. CD3^+^CD4^+^, CD3^+^CD8^+^ and CD11c^+^CD83^+^ double positive cells, as well as CD3^+^, CD3^−^CD8^+^, CD20^+^, CD11c^+^, CD11c^−^ DC-SIGN^+^ and PM-2K^+^ single positive cells were automatically counted in the chorion areas, while CD20^−^MamuLa-DR^+^ single positive cells were automatically counted in both the chorion and the epithelium zones. Results were expressed as the number of cells per mm^2^ of chorion or epithelium.

Chemokine^+^ surfaces were automatically quantified in the manually defined zones of chorion and epithelium. Results were expressed as the sum of simply stained surfaces/total surface analyzed. For each staining, at least 1.2 mm² of chorion and 0.9 mm² of epithelium surfaces were analyzed per macaque. Quantifications were performed on 5 to 8 macaques, 2 to 3 biopsies per macaque amongst 4 biopsies sampled both at D-30 and 48 hours post-rs-IL-7gly, and 3-4 or 2-3 independent zones of chorion or epithelium were defined, respectively (except for CD11c^+^CD83^+^ cells quantification: 4 macaques). Image analysis quantification was performed independently by at least two people.

Lymphoid follicles were enumerated in high-powered entire reconstitutions of vaginal sections acquired with the Lamina™ Multilabel Slide Scanner (PerkinElmer, Courtaboeuf, France), manually counting 8 to 14 sections per macaque. The chorion surfaces were manually outlined, and lymphoid follicles were highlighted manually and then automatically quantified with CaseViewer software (3DHISTECH, Budapest, Hungary). The results were expressed as the number of follicles per 50 mm^2^ of chorion. Two people performed image analysis quantifications independently.

The proportions of T- and B-cells in lymphoid follicles were assessed on vaginal mucosa at necropsy, using ImageJ (1.52p version) on sections acquired with a 20x objective. Lymphoid follicles were manually defined on the DAPI channel, manual thresholds were applied on CD3^+^ and CD20^+^ stains to automatically measure CD3^+^ and CD20^+^ stained surfaces. Nuclei (DAPI) were also quantified in the follicles. Results were expressed as the number of B-cells over total cell numbers in each follicle. The quantifications were performed on 7 to 21 follicles per macaque. Two people performed image analysis quantifications independently.

### Quantification of total and DT-specific IgGs and IgAs by enzyme-linked immunosorbent assay

Total and DT-specific immunoglobulin (IgGs and IgAs) were quantified in CVLs using in-house ELISA, as described in the supplementary material. Total IgA or IgG concentrations were determined by interpolation, using the calibration line of IgA or IgG standards, respectively. For the quantification of specific Igs, samples with a signal at least twice above the background were considered positive. The results were expressed as OD (in IgA or IgG anti-DT ELISA) over the concentration of IgAs or IgGs in a given sample.

### Preparation of cells for ELISPOT assay

The lower FGT and the iliac LN cells were isolated as described in the supplementary material. Cell number and viability were determined by trypan blue exclusion.

### Quantification of antibody-secreting cells by B-cells ELISPOT

Antibody secreting cells (ASCs) were assayed in Multiscreen HA plates (Merck Milipore, Molsheim, France) coated with DT (10μg/mL), as described in the supplementary material. The spot numbers were reported as DT-specific ASCs per million PBMCs. In the wells used as specific controls, the ELISPOT Reader detected 0 to 2 spots for the anti-IgAs, and 1 to 4 spots for the anti-IgGs, in DT-coated wells with cells from non-immunized animals, uncoated wells with cells from immunized animals or wells incubated without cells.

### Statistical analysis

Non-parametric Mann-Whitney U tests, Wilcoxon Signed-Rank Tests and multivariate analysis of variance (MANOVA) with the post hoc analyses, and Fisher least significant difference (LSD) tests were performed using StatView (5.0 version, Abacus software). A p value <0.05 is considered as significant.

## RESULTS

### Local administration of rs-IL-7gly elicits chemokine expressions by the vaginal mucosa

In a first series of experiments, recombinant glycosylated simian IL-7 (rs-IL-7gly) was injected into the vaginal mucosa of 8 healthy rhesus macaques (1 to 10 ng/injection site, 4 injections per animal, together with Indian ink in order to locate the injection sites). Twenty-four and forty-eight hours after inoculation, IL-7-injected and control zones were biopsied and mRNAs coding for 12 chemokines were quantified by qRT-PCR. Six chemokines (CCL5, CCL19, CCL28, CXCL8, CXCL10 and CXCL12) demonstrated a significantly higher transcription level in the IL-7-treated zones sampled 48 hours after inoculation, compared to control biopsies (n=3 and n=9 macaques, respectively; **Figure 1A**). In contrast, these overexpressions were neither observed in biopsies sampled 24 hours after rs-IL-7gly injection (n=5 macaques) nor in biopsies injected with Indian ink alone (n=8 macaques), with the exception of CXCL10 (**Figure 1A**), suggesting a consequence of the needle prick itself. By analyzing the expression of chemokines in microdissected epithelium and chorion, we then demonstrated that IL-7 driven chemokine transcription was located in the chorion (CCL19 and CXCL12) or in the epithelium (CCL28) or in both (CCL5, CXCL8 and CXCL10; **Figure 1B**). These data demonstrate that a few nanograms of rs-IL-7gly, directly injected into the vaginal mucosa, are sufficient to trigger a significant enhancement of local chemokine expressions, which can be measured 48 hours after administration.

**Figure 1.**
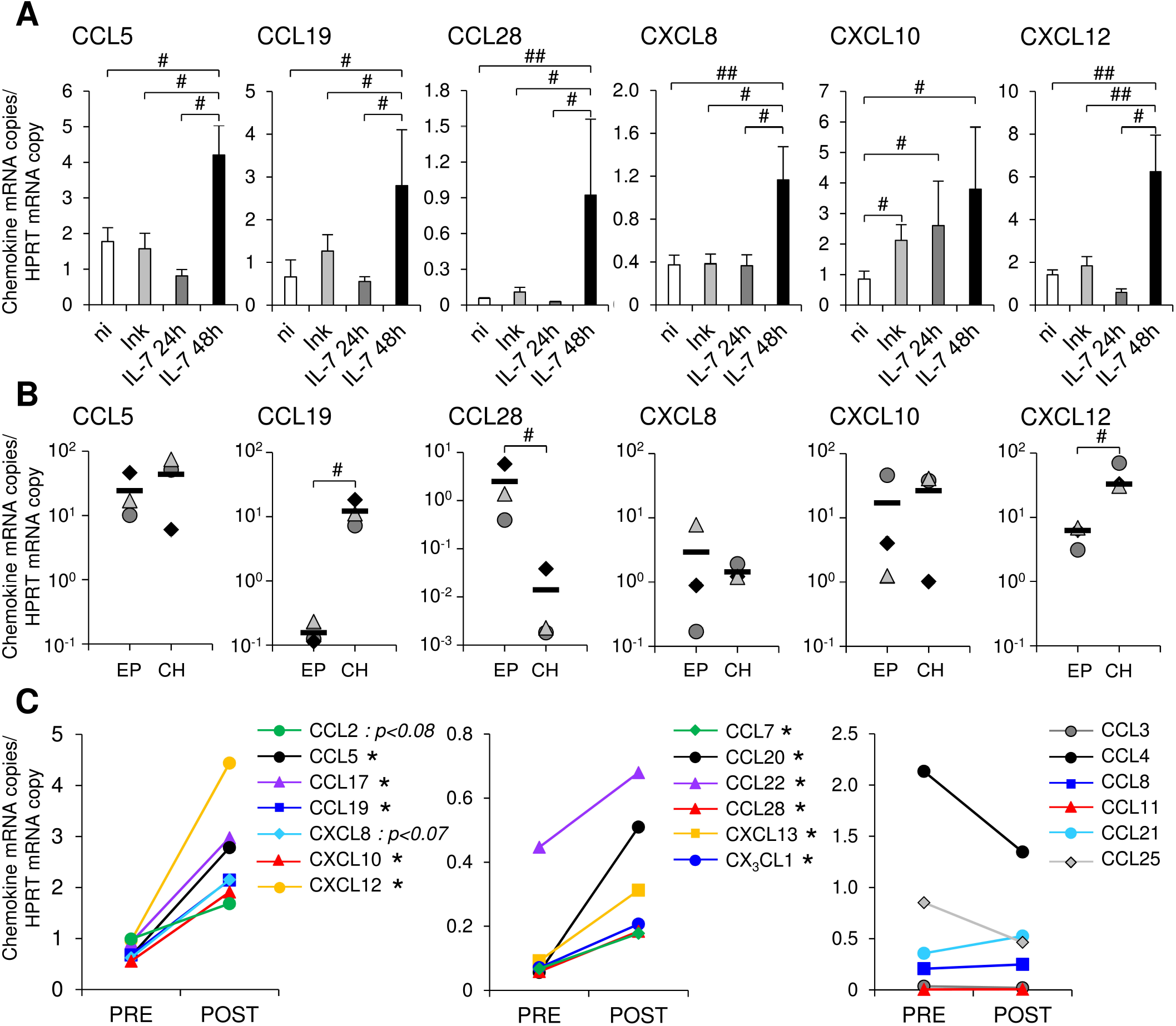
Topical administration of rs-IL-7gly induces local chemokine transcription in the vaginal mucosa. **(A)** mRNAs coding for CCL5, CCL19, CCL28, CXCL8, CXCL10 and CXCL12 were quantified in vaginal biopsies (2-4 biopsies per macaque) sampled 24 hours or 48 hours after rs-IL-7gly-injection (IL-7 24h, dark gray bars, n=5 macaques; IL-7 48h, black bars, n=3 macaques), 24 and 48 hours after injection with Indian ink alone (Ink, light gray bars, n=8 macaques) and from non-injected healthy rhesus macaques (ni, white bars, n=9). Data were normalized to HPRT mRNAs simultaneously quantified together with the chemokines (Chemokine mRNA copies/HPRT mRNA copy). Bars and error bars represent means and SEM, respectively. ##: p<0.01, #: 0.01<p<0.05 (one-tailed Mann-Whitney U test). **(B)** mRNAs coding for CCL5, CCL19, CCL28, CXCL8, CXCL10 and CXCL12 were quantified in pluristratified epithelium (EP) or chorion (CH) microdissected from vaginal biopsies sampled 48 hours after rs-IL-7gly administration (n=3 macaques). Each symbol represents one macaque (6-9 microdissected zones per macaque), and horizontal black bars represent means. #: p<0.05 (Mann-Whitney U test). **(C)** mRNAs coding for 19 chemokines were quantified in vaginal biopsies (4 biopsies per macaque) sampled from macaques one month before (PRE, n=5) and 48^H^ after the administration of 10μg (n=3) or 15μg (n=2) of rs-IL-7gly (POST), by vaginal spray. Data were normalized to HPRT mRNAs simultaneously quantified together with the chemokines (Chemokine mRNA copies/HPRT mRNA copy). Each point represents the mean value obtained for the 5 macaques at each time point. *: p<0.05 (Wilcoxon Signed-Rank Test).

We then investigated the effect of the non-traumatic administration of rs-IL-7gly directly sprayed onto the vaginal mucosal surface of healthy rhesus macaques. Nine healthy macaques were administered with 1μg (n=2), 5μg (n=2), 10μg (n=3) or 15μg (n=2) of rs-IL-7gly. The expression of 19 chemokines and 5 cytokines was measured by qRT-PCR in vaginal biopsies sampled 48 hours after IL-7 administration. Interestingly, while the administration of 1 and 5μg did not impact the mucosal expression of chemokines, animals treated with either 10μg or 15μg of rs-IL-7gly demonstrated a significantly enhanced mRNA level for 11 chemokines (**Figure 1C** and **Supplementary Figure 1**) compared to the baseline. Among these, CCL5, CCL17 (TARC), CCL19, CXCL10 and CXCL12 were constitutively expressed at the baseline (**Figure 1C**, left panel), while CCL7, CCL20, CCL22 (MDC), CCL28, CXCL13 (BCA-1), CX_3_CL1 (Fractalkine) were scarcely transcribed before IL-7 treatment (**Figure 1C**, central panel). Finally, the vaginal expression of CCL3, CCL4, CCL8 (MCP-2), CCL11 (Eotaxin), CCL21 and CCL25 was not significantly modified upon stimulation with IL-7 (**Figure 1C**, right panel). Among the cytokines tested, only IL-17A and TSLP demonstrated enhanced expression in IL-7-treated vaginal mucosa (mean IL-17A expression: 0.04 and 0.32 copies/HPRT copy in baseline and D2 samples, respectively; p<0.01; mean TSLP expression: 1.01 and 4.61 copies/HPRT copy in baseline and D2 samples, respectively; p<0.01; n=5 monkeys; **Supplementary Figure 2**).

We then analyzed the expression of chemokines, at the protein level, in IL-7-treated vaginal tissue samples taken 30 days before and 48 hours after the administration of rs-IL-7gly (10μg by spray). Increased amounts of these chemokines were observed by immunochemistry in either the epithelium or the chorion in samples gathered 48 hours after rs-IL-7gly administration (**Figure 2A**), confirming mRNA quantifications. Quantification of labeled surfaces using ImageJ software (see Methods) demonstrated that CCL7, CCL2, CXCL13 and CCL5 expressions were increased in the chorion (1.5-, 1.9-, 2- and 3.4-fold, respectively; p<0.02; **Figure 2B**) while only CCL7, CCL2 and CCL5 were overexpressed in the epithelium (1.7-, 1.8- and 1.9-fold, respectively; p<0.05; **Figure 2C**).

**Figure 2.**
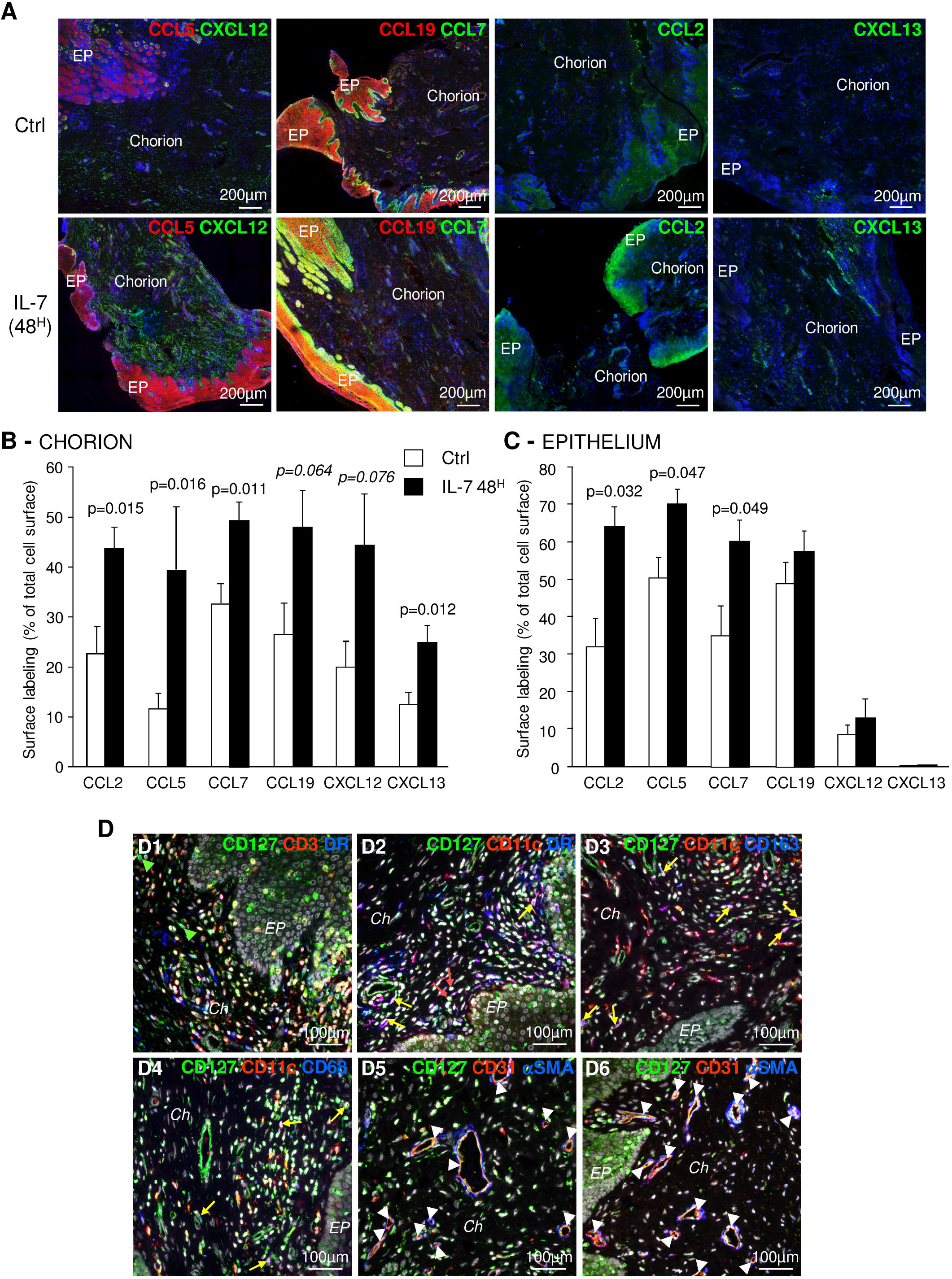
Topical administration of rs-IL-7gly increases local chemokine expression in the vaginal mucosa. **(A)** Sections of vaginal mucosa biopsies sampled 30 days before (Ctrl), or 2 days after (IL-7 48^H^) the administration of 10μg of rs-IL-7gly by vaginal spray were immunostained with anti-CCL5 or -CCL19 (red) antibodies, in combination with anti-CXCL12 or -CCL7 (green) antibodies, and anti-CCL2 or −CXCL13 (green) antibodies. Nuclei were stained with DAPI (blue). *EP: Pluristratified Epithelium.* **(B, C)** The expression of CCL2, CCL5, CCL7, CCL19, CXCL12 and CXCL13 was quantified by image analysis of immunohistofluorescent staining. Data are expressed as percentages of total chorion **(B)** or the epithelium **(C)** surface labeled by the different antibodies. Each bar represents the mean ± SEM of quantifications performed on 5-8 macaques (2-3 biopsies per animal) sampled 30 days before (Ctrl, white bars) and 48 hours after (IL-7 48^H^, black bars) the administration of 10μg of rs-IL-7gly. Statistical significance of the differences between IL-7 treated and control animals are shown at the top of the figure (Mann-Whitney U test). **(D)** Sections of vaginal mucosa were labeled with anti-CD127 and combinations of anti-CD3, anti-MamuLa-DR, anti-CD11c, anti-CD163, anti-CD68, anti-CD31 and anti-αSMA antibodies. Nuclei were stained with DAPI (grey). *Green arrowheads identify CD127^+^CD3^−^MamuLa-DR^−^ cells; Red arrows identify CD127^+^CD11c^+^MamuLa-DR^−^ cells; Yellow arrows identify: CD127^+^CD11c^+^MamuLa-DR^+^ (D2), CD127^+^CD11c^+^CD163^+^ (D3), or CD127^+^CD11c^+^CD68^+^ (D4) triple positive cells; White arrowheads identify CD127^+^CD31^+^ endothelial cells. EP: Pluristratified Epithelium; Ch: Chorion; DR: MHC-II MamuLa-DR.*

These different chemokines are often described as produced either by myeloid cells or by resident mucosal cells, suggesting that these cells could sense IL-7 through expression of the IL-7 receptor. Accordingly, we investigated the expression of CD127 (the alpha chain of the IL-7 receptor) by various types of cells composing the vaginal mucosa. In addition to CD3^+^ T-cells, many different CD3^−^MamuLa-DR^+^ cells effectively expressed CD127 (**Figure 2D** and **Supplementary Figure 3**). These cells were mostly CD11c^+^ dendritic cells (**Figure 2D**, panel D2; yellow arrows). Likewise, some CD11c^+^CD68^+^ or CD11c^+^CD163^+^ cells, likely representing mucosal pro-inflammatory “M1” macrophages or cells with a mixed “M1/M2” phenotype, also expressed CD127 (**Figure 2D**, panels D3 and D4; yellow arrows). Finally, a few CD11c^+^MamuLa-DR^−^ cells, presumably NK-cells, also expressed CD127 (**Figure 2D**, panel D2; red arrows).

Furthermore, CD127 was expressed by CD31^+^ endothelial cells but not by surrounding αSMA^+^ (alpha-smooth muscle actin) cells (**Figure 2D**, panels D5 and D6; white arrowheads). In contrast, some isolated αSMA^+^ cells expressed CD127. Finally, epithelial cells also presented CD127 staining (**Figure 2D**, panels D1, D2 and D6 and **Supplementary Figure 3**, panels B, C, D), with a more distinct expression by basal epithelial cells. Interestingly, we also evidenced both CD127 and CD132 transcription in epithelial cells isolated from the vaginal mucosa of healthy macaques, confirming the expression of the entire IL-7 receptor (**Supplementary Figure 4**). Nonetheless, the expression of CD127 by epithelial cells remained significantly lower than on T-cells isolated from blood or secondary lymphoid organs (SLO) (7.8-fold and 4.2-fold in blood and SLO T-cells, respectively; p<0.05; **Supplementary Figure 4**).

Therefore, many cell types that compose the vaginal mucosa express the IL-7 receptor and may contribute to the chemokine production observed in IL-7-treated vaginal mucosa.

Altogether, these results demonstrate that the non-traumatic administration of rs-IL-7gly at the surface of the vaginal mucosa stimulates the local expression of a set of chemokines which may trigger immune cell migrations to the IL-7-treated mucosa.

### Vaginal administration of IL-7 triggers the recruitment of immune cells into the vaginal chorion

We further evaluated the consequences of IL-7 mucosal treatment on the distribution of immune cells within the vaginal mucosa by performing immunohistofluorescent staining on both biopsies 2 days after administration of rs-IL-7gly by spray (10μg/animal, n=8 macaques) and control biopsies. A significant increase in cell density, evidenced by nuclei counts per unit of tissue surface, characterized the mucosa treated with IL-7 (1994±168 and 3518±326 cells per mm^2^ of chorion in control and IL-7-treated samples, respectively; p=0038; data not shown). CD4^+^ and CD8^+^ T-cells, NK cells, B-cells, myeloid DCs (mDCs), macrophages, and MamuLa-DR^+^ APCs were further quantified in the chorion on whole tissue sections using ImageJ software (**Figure 3A**). These quantifications confirmed that local rs-IL-7gly administration triggers massive infiltration of all the immune cell subsets we searched for in the vaginal mucosa (5.4-, 5.7-, 3.4- and 3.5-fold increase over baseline values for CD4^+^ T-cells, CD8^+^ T-cells, B-cells and NK cells, respectively; p<0.01, <0.01, <0.05 and <0.01; **Figure 3B**). Similarly, APC numbers (DR^+^CD20^−^ cells) were also massively increased in the vaginal chorion following local administration of rs-IL-7gly (2.9-fold increase; p<0.05). Among these, we identified CD11c^+^ mDC, DC-SIGN^+^ macrophages and PM-2K^+^ tissue macrophages (4.7-, 3.1- and 1.6-fold increase; p<0.01, p<0.05 and p=0.076, respectively; **Figure 3C**). We noted that most of the PM-2K^+^ tissue macrophages also expressed DC-SIGN in the vaginal mucosa (data not shown). Moreover, after the administration of rs-IL-7gly, CD11c^+^DC-SIGN^+^ mDCs concentrated underneath the epithelium (**Figure 3A**) and expressed CD83 (21.4-fold increase after rs-IL-7gly treatment; p=0.021; **Figures 3D-E**). Furthermore, following administration of rs-IL-7gly by spray, a significant increase in the numbers of APCs expressing MamuLa-DR was observed in the vaginal epithelium (39±5 and 70±8 cells/mm^2^ in control and IL-7-treated macaques, respectively, p=0.013; **Figures 3G-H**).

**Figure 3.**
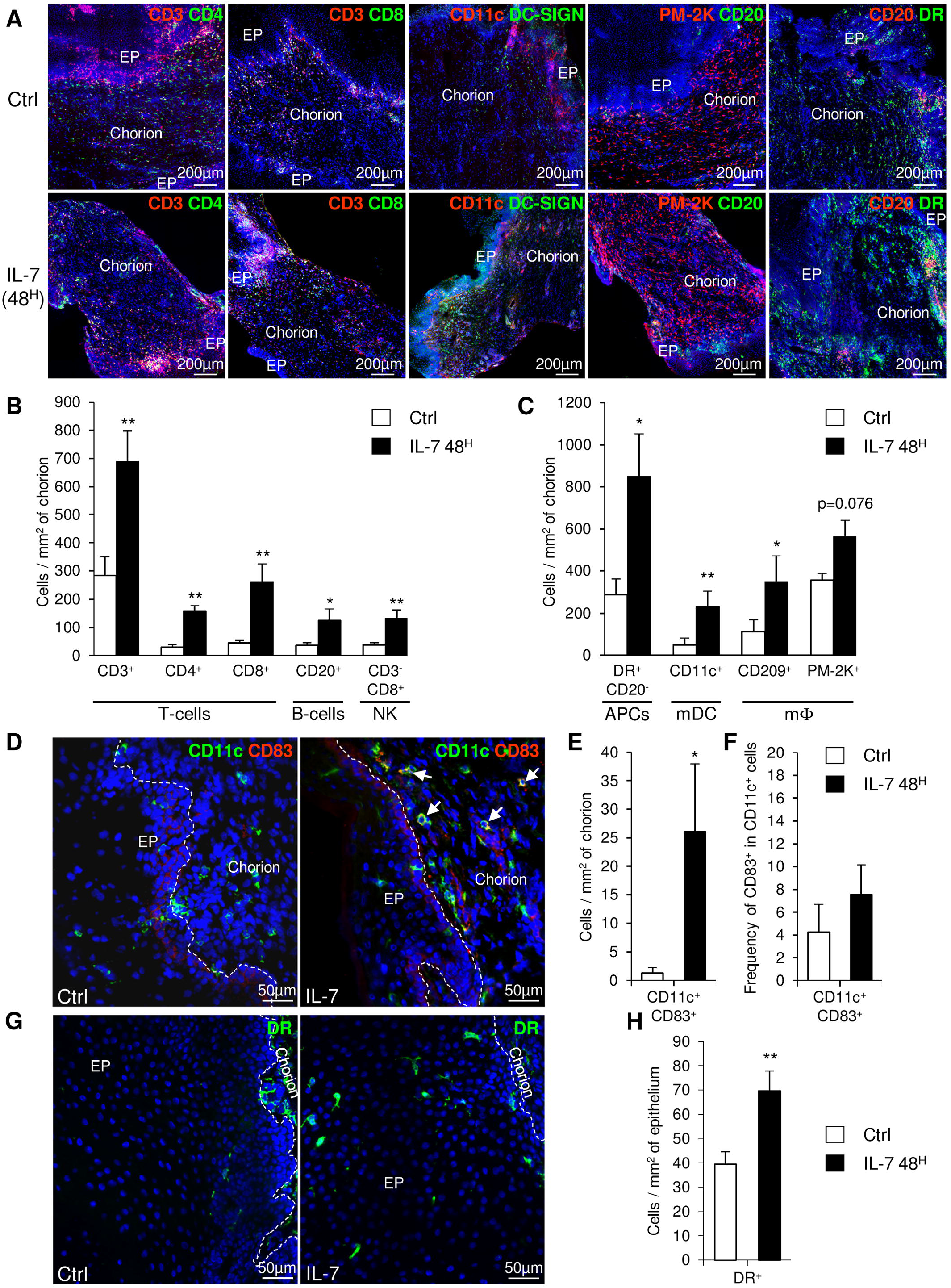
Topical administration of rs-IL-7gly induces the recruitment of immune cells into the vaginal chorion. **(A)** Sections of vaginal mucosa biopsies sampled 30 days before (Ctrl), or 48 hours after (IL-7 48^H^) the administration of 10μg of rs-IL-7gly by vaginal spray were labeled with anti-CD3, -CD11c, -PM-2K and -CD20 antibodies, in combination with anti-CD4, -CD8, -DC-SIGN, -CD20 or -MHC-II MamuLa-DR antibodies. Nuclei were stained with DAPI (blue). **(B, C)** Cell infiltration was quantified by image analysis of immunohistofluorescent staining and expressed as numbers of cells per mm^2^ of chorion. Each bar represents the mean ± SEM of quantifications performed on 5-8 macaques (2-3 biopsies per animal) sampled 30 days before (Ctrl, white bars) and 48 hours after (IL-7 48^H^, black bars) the administration of 10μg of rs-IL-7gly. **(D)** Sections of vaginal mucosa biopsies sampled 30 days before (Ctrl), or 48 hours after (IL-7) the administration of 10μg of rs-IL-7gly by vaginal spray were labeled with anti-CD11c (green) and anti-CD83 (red) antibodies. Nuclei were stained with DAPI (blue). Arrows identify CD11c^+^CD83^+^ mature myeloid dendritic cells. CD11c^+^CD83^+^ cells were quantified by image analysis of immunohistofluorescent staining on vaginal mucosa biopsies sampled from macaques (n=4) 30 days before (Ctrl, white bars) and 48 hours after (IL-7 48^H^, black bars) the administration of 10μg of rs-IL-7gly and expressed as number of double positive cells per mm^2^ of chorion ± SEM **(E)** and as the frequency of CD83^+^ cells in CD11c^+^ cells **(F)**. **(G)** Sections of vaginal mucosa biopsies sampled 30 days before (Ctrl), or 48 hours after (IL-7) the administration of 10μg of rs-IL-7gly by vaginal spray were immunostained with anti-MamuLa-DR antibodies (green). Nuclei were stained with DAPI (blue). **(H)** MamuLa-DR^+^ cells were quantified by image analysis of immunohistofluorescent staining on vaginal mucosa biopsies sampled from macaques (n=7) 30 days before (Ctrl, white bars) and 48 hours after (IL-7 48^H^, black bars) the administration of 10μg of rs-IL-7gly. **: p<0.01, *: 0.01<p<0.05 (Mann-Whitney U test). *EP: Pluristratified Epithelium; DR: MHC-II MamuLa-DR.*

These data demonstrate that, in the vaginal mucosa, the chemokine expressions induced by IL-7 treatment trigger the migration of APCs, B-cells, T-cells and NK cells into the chorion, and lead to the activation of mDCs, a prerequisite to the development of immune responses to local antigenic stimulation. Besides, the large numbers of MamuLa-DR^+^ APCs localized in the epithelium after rs-IL-7gly administration could certainly help subsequently administered immunogens penetrate the mucosa.

### IL-7-adjuvanted vaginal vaccine stimulates strong mucosal antibody responses

We then tested the capacity of IL-7 to serve as an adjuvant in a mucosal immunization protocol against a model antigen. Six female rhesus macaques were immunized against diphtheria toxoid (DT) through local administration of the antigen (7μg of DT in 200μL of PBS) sprayed directly into the vaginal lumen. Two days prior to the immunization, these animals had been treated by local administration of rs-IL-7gly (group IL-7+DT; n=3; 10μg) or PBS (group PBS+DT; n=3), by the same route. Anti-DT antibodies were quantified in mucosal secretions sampled over a 15-week period by cervico-vaginal lavages (**Figures 4A-B**). Interestingly, DT-specific IgGs were detected in the vaginal secretions of all IL-7-treated DT-immunized animals by week 2 or 3 (W2/3). These antibodies remained at a higher concentration for the subsequent 15 weeks, compared to animals receiving DT without pretreatment with IL-7 (**Figure 4A**; p<0.001). Indeed, among the control animals, one never developed any detectable DT-specific IgG response and the others showed a weak and sporadic response by W4. Similarly, DT-specific IgA responses appeared earlier and were stronger in the IL-7-treated DT-immunized animals as compared to the low and sporadic IgA response being detectable by W4/5 in 2 of the animals immunized without IL-7 treatment (**Figure 4B**; p=0.029).

**Figure 4.**
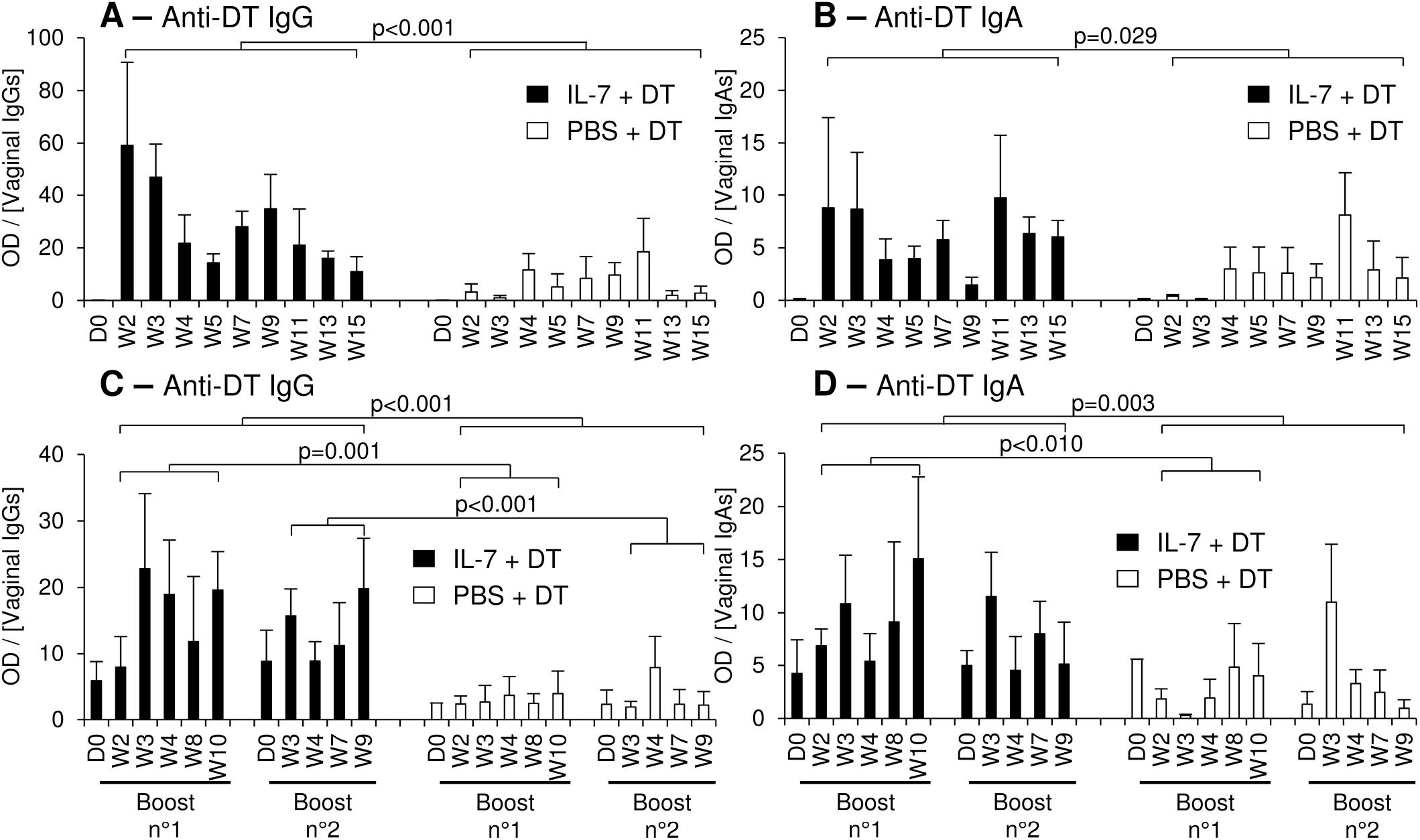
Topical administration of DT leads to a stronger mucosal immune response after local administration of rs-IL-7gly. Specific anti-DT IgGs **(A, C)** and IgAs **(B, D)**, were quantified by ELISA in vaginal secretions of 6 rhesus macaques that received vaginal administration of either 10μg of rs-IL-7gly (black bars; n=3) or PBS (white bars, n=3), followed, at day 2 (D2), by local administration of Diphtheria Toxoid (DT). Two boosts were performed at 16 and 31 weeks following prime immunization, using the same protocol. All administrations were performed by vaginal spray. Specific anti-DT antibody responses are expressed as optical density over IgG or IgA concentration in each CVL sample. Bars and error bars represent means and SEM at any time-point for the 3 animals from each group. Samples containing blood contaminations due to menstruations were excluded. Statistical differences between IL-7-treated and untreated monkeys are shown (MANOVA Test). *D0: Administration of rs-IL-7gly or PBS; D2: Administration of DT; W: Week post-DT administration.*

Boost immunizations were performed 16 and 31 weeks after prime immunization, using the same protocol. In all three IL-7-treated DT-immunized macaques, a rebound in both IgG and IgA vaginal DT-specific responses was observed, despite the fact that IgG response did not reach the levels observed early after prime immunization (**Figures 4C-D**). In contrast, no significant increase of the vaginal antibody responses (IgG or IgA) was observed in the animals immunized without IL-7 pretreatment. In these animals, sporadic DT-specific IgGs and IgAs were detected, their concentrations remaining lower than those in IL-7-treated animals (**Figures 4C-D**; p<0.001 and p=0.003, respectively, for both boosts).

Immunoglobulins in vaginal secretions can be either produced by resident antibody-secreting cells (ASCs) in the mucosa or excreted by transudation of serum antibodies. We thus looked for DT-specific ASCs in the vaginal mucosa and draining LNs sampled from animals sacrificed 2 weeks after a third boost immunization performed 24 weeks after the second boost.

In both PBS- and IL-7-treated DT-immunized macaques, DT-specific IgA^+^ plasma cells outnumbered DT-specific IgG^+^ plasma cells. However, a higher density of DT-specific IgA^+^ plasma cells characterized both the vaginal walls and the fornix (i.e. the glandular-rich mucosal region around the uterine cervix) of the IL-7-treated DT-immunized macaques (**Figure 5A**). Similarly, we evidenced a higher density of anti-DT ASC in the iliac LNs of animals treated with IL-7. However, in LNs, IgG^+^ plasma cells predominate among the anti-DT ASCs (**Figure 5B**).

**Figure 5.**
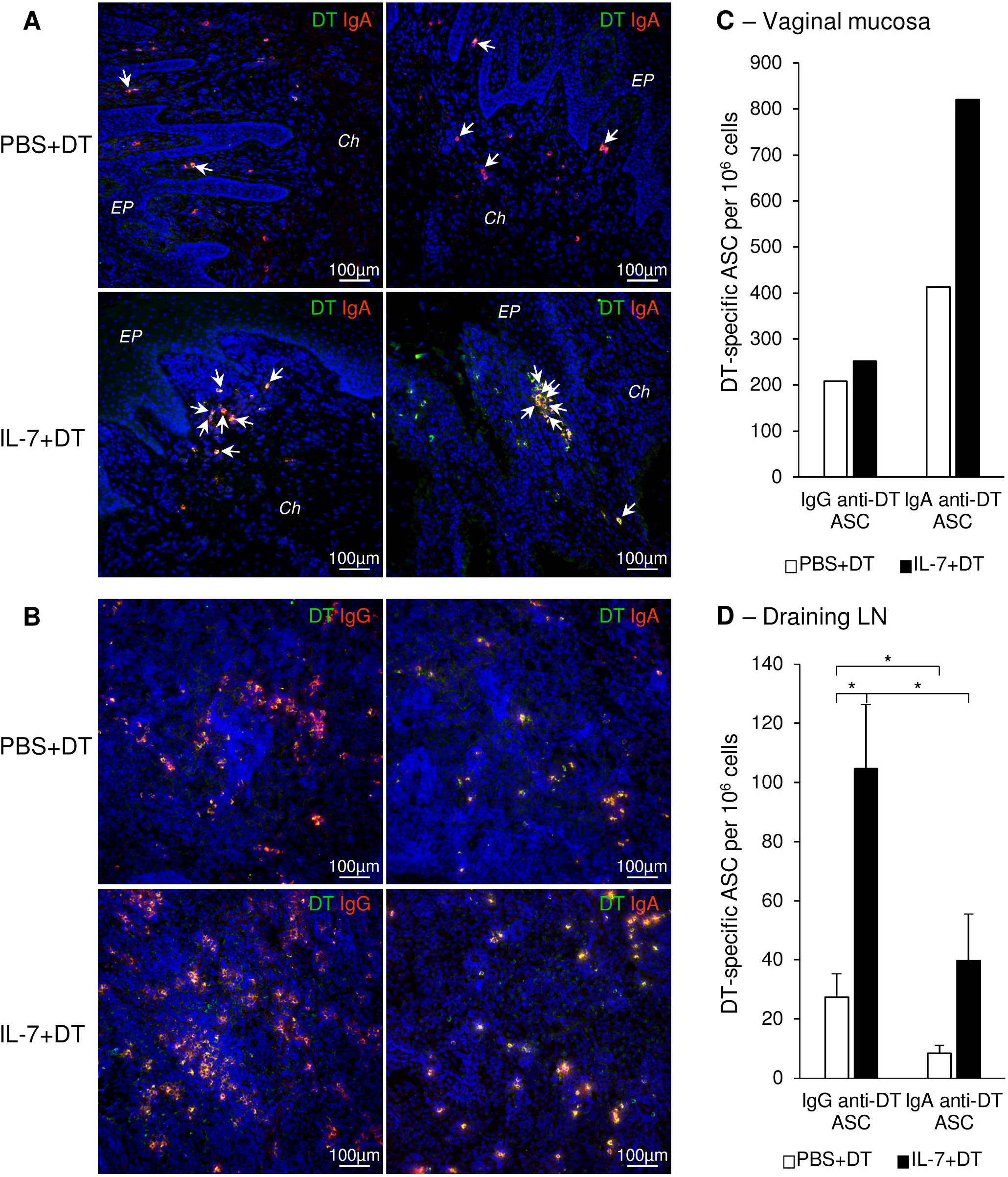
Preferential localization of DT specific IgAs plasma cells in the vaginal mucosa following rs-IL-7gly-adjuvanted mucosal immunization. Sections of vaginal mucosa **(A)**, or iliac lymph nodes **(B)**, sampled at necropsy (i.e. 2 weeks after the fourth mucosal immunization) from PBS+DT (top panels) and IL-7+DT (bottom panels) -immunized macaques, were incubated with DT and immunostained with anti-DT antibodies (green) and either anti-IgA or anti-IgG (red) antibodies to reveal IgA and IgG anti-DT plasma cells, respectively. Nuclei were stained with DAPI (blue). Representative examples of the upper part of the vagina (A, left panels) and vaginal fornix (A, right panels) or of the draining lymph nodes (B) are shown. DT-specific plasma cells are yellow (A, B) and arrows indicate DT-specific IgA plasma cells in vaginal mucosa (A). *EP: Pluristratified Epithelium; Ch: Chorion.* **(C, D)** IgG- and IgA-producing DT-specific plasma cells (ASC) were quantified by B-cell ELISPOT on isolated cells from the vaginal chorion of macaques immunized with PBS+DT (White bars, n=1) or IL-7+DT (Black bars, n=1), sampled at necropsy **(C)**, and on isolated cells from iliac lymph nodes from macaques immunized with PBS+DT (White bars, n=3) or IL-7+DT (Black bars, n=3), sampled at necropsy **(D)**. Results are expressed as IgG or IgA anti-DT-specific plasma cells per 10^6^ cells. Bars and error bars represent means and SEM, respectively (two independent experiments performed in duplicate; *: p<0.05 (Mann-Whitney U test)). *LN: Lymph nodes; ASC: antibody secreting cells.*

These data were confirmed by the quantification of IgA^+^ anti-DT ASCs by ELISPOT performed on purified immune cells either from the vaginal mucosa (**Figure 5C**) or the iliac LNs of macaques from each group (IgG ASC: 81, 148 and 85 spots/10^6^ cells and 17, 43 and 22 spots/10^6^ in macaques from the IL-7+DT and the PBS+DT groups, respectively; p<0.05. IgA^+^ DT-specific ASCs: 55, 56 and 8 spots/10^6^ cells as compared to 10, 12 and 3 spots/10^6^ cells in macaques from the IL-7+DT and the PBS+DT groups, respectively; **Figure 5D**).

### IL-7-adjuvanted vaginal vaccine allows stronger systemic immune responses

Having demonstrated the adjuvant potential of rs-IL-7gly through its capacity to improve mucosal DT-specific antibody responses, we further analyzed B-cell responses in both secondary lymphoid organs and blood.

The frequency of DT-specific ASCs of IgG and IgA isotypes was determined in blood samples collected at different time points after prime immunization and following the different boosts.

Two weeks after primary immunization, higher frequencies of DT-specific IgG^+^ ASCs were observed in macaques immunized by local administration of IL-7+DT compared to those receiving DT immunization alone (W2: 193, 82 and 73 DT-specific IgG^+^ ASC/10^6^ PBMCs in the IL-7-treated macaques compared to 2 and 64 in the control macaques) (**Figure 6A**). At later time points, these frequencies remained higher in IL-7-treated macaques (55, 41 and 130 DT-specific IgG^+^ ASC/10^6^ PBMCs at W3 and 54, 29 and 89 at W5 in the 3 IL-7-treated macaques as compared to 11, 31, 13 and 7, 27, 39 in control animals; **Figure 6A**). The frequency of DT-specific IgG^+^ ASCs increased after boost immunizations in both groups of macaques; the rebound of the immune response being higher following the second boost in IL-7-treated macaques. Unlike IgG^+^ ASCs, circulating DT-specific IgA^+^ ASCs remained low throughout the immunization protocol, their frequencies being slightly higher in IL-7-treated DT-immunized macaques (**Figure 6B**).

**Figure 6.**
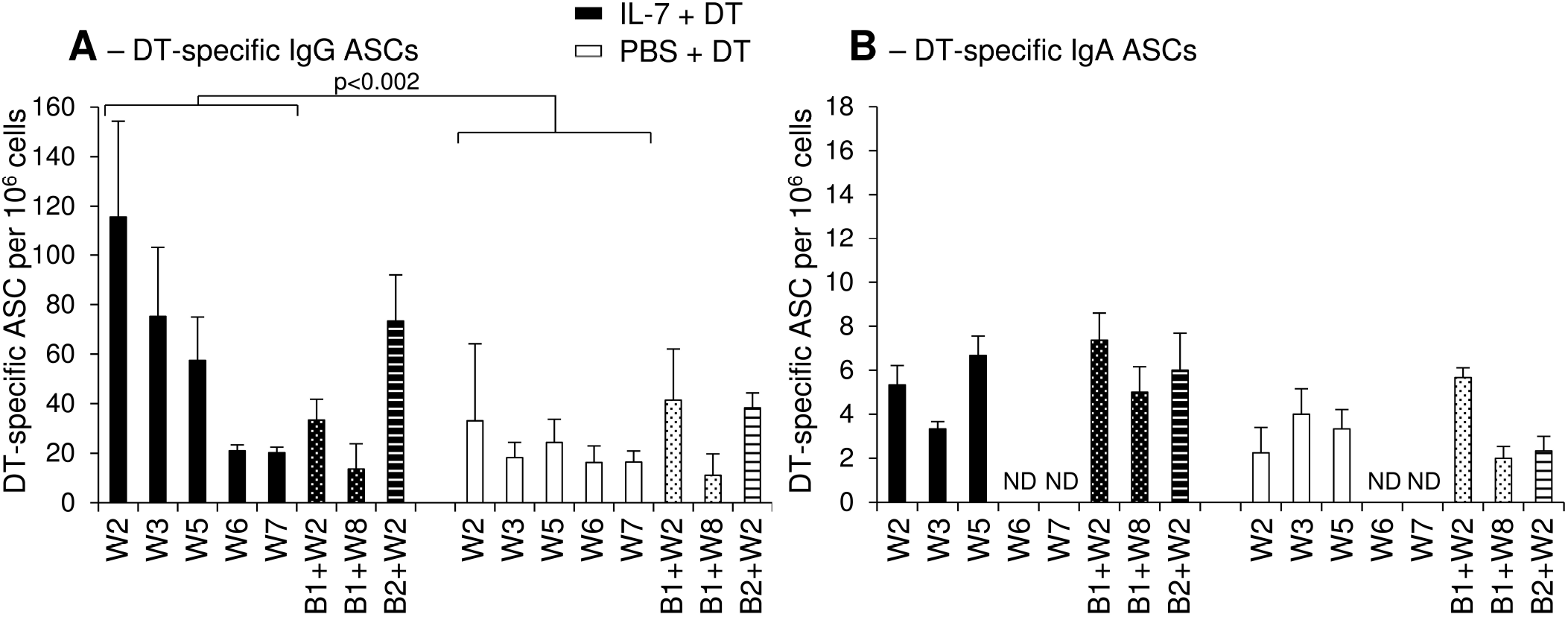
Increased numbers of circulating DT-specific IgG antibody-secreting cells after rs-IL-7gly-adjuvanted vaginal immunization. DT-specific ASC of IgG **(A)** and IgA **(B)** isotypes were quantified by B-cell ELISPOT on peripheral blood mononuclear cells (PBMCs) from PBS+DT (White bars, n=3) and IL-7+DT (Black bars, n=3) immunized macaques sampled after prime immunization (plain bars) or after boost #1 (dotted bars) or boost #2 (hatched bars). Results are expressed as the number of IgG or IgA anti-DT-specific cells per 10^6^ PBMC. Bars and error bars represent means and SEM obtained in two independent experiments performed in duplicate. Statistical differences between IL-7-treated and PBS-treated immunized monkeys are shown (MANOVA Test). *ND: Not determined.*

Altogether, these data demonstrate that rs-IL-7gly acts as a mucosal vaccine adjuvant. Its administration at the mucosal surface prior to immunization, accelerates, enhances and stabilizes the mucosal antigen-specific antibody responses triggered by local antigenic stimulation.

### IL-7-adjuvanted mucosal immunization induces ectopic lymphoid follicles in vaginal mucosa

To further explore the mechanisms involved in the induction of mucosal immunity in the vaginal mucosa of macaques treated with IL-7, we quantified the expression of chemokines involved in the development of tertiary lymphoid structures (TLS) in vaginal biopsies sampled at necropsy. The amount of mRNA encoding CCL19, CCL21, CXCL12, and CXCL13 (chemokines known to trigger lymphocytes trafficking and aggregation in tissues) was increased in vaginal tissues collected from IL-7-adjuvanted immunized macaques (6.1-, 4.9-, 25.8- and 54.2-fold over pre-immunization values for CCL19, CCL21, CXCL12 and CXCL13, respectively; p<0.05 as compared to control animals; **Figure 7A**). Similar data were obtained in biopsies sampled 4 weeks after each rs-IL-7gly administration during the immunization protocol (data not shown).

**Figure 7.**
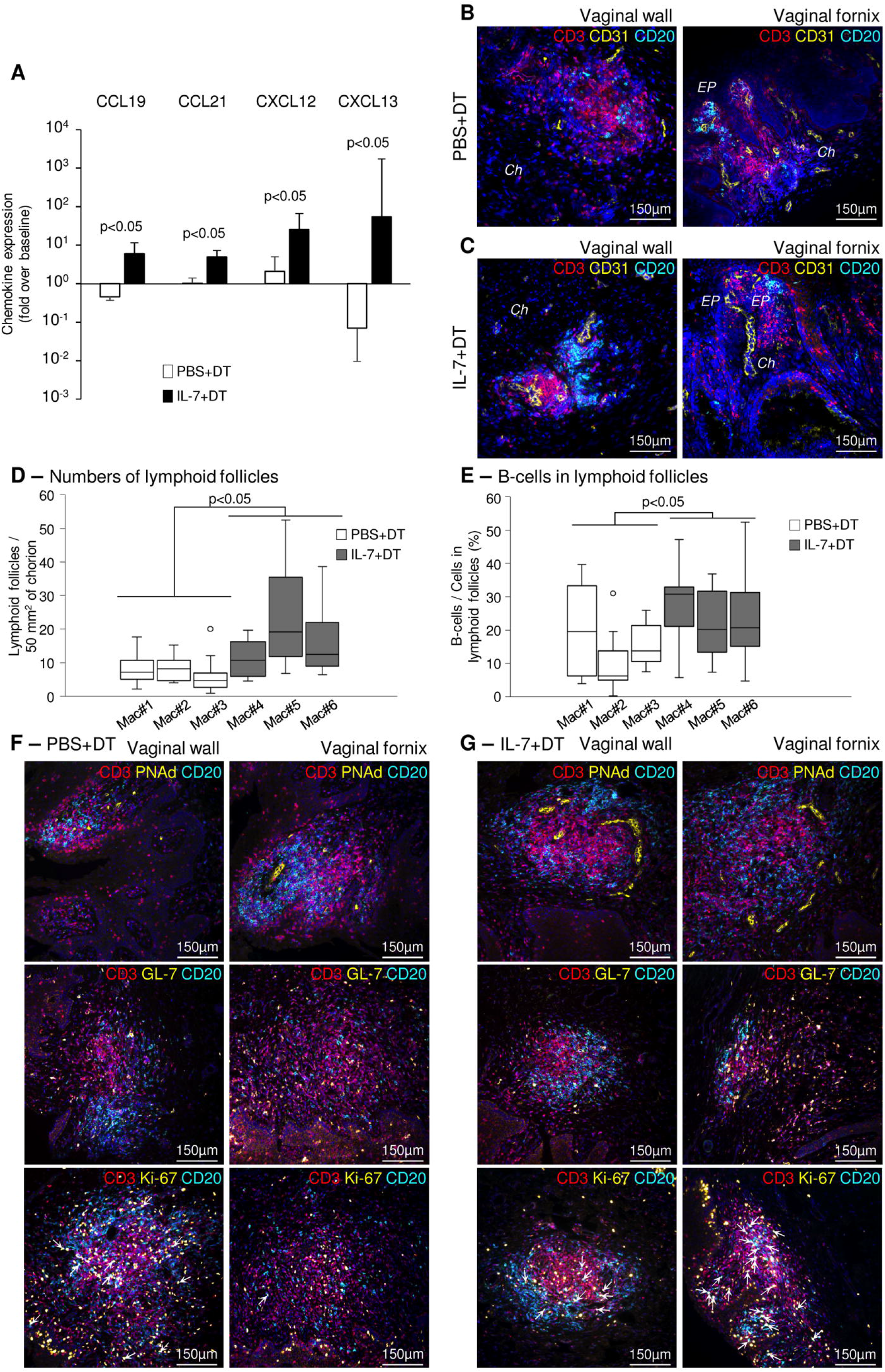
Induction of ectopic lymphoid follicles in the vaginal mucosa of IL-7-treated macaques. **(A)** mRNAs coding for CCL19, CCL21, CXCL12 and CXCL13 were quantified in vaginal biopsies (n=3-4 per macaque) sampled at baseline and at necropsy from IL-7+DT (black bars, n=3) and PBS+DT (white bars, n=3) immunized macaques. Data are presented normalized to HPRT mRNAs simultaneously quantified together with the chemokines (chemokine mRNA copies/HPRT mRNA copy). Bar and error bars represent the fold increase over baseline values and SD. Statistical differences between IL-7-treated and PBS-treated monkeys are shown (Mann-Whitney U test). **(B, C)** Sections of vaginal walls (left panels) and vaginal fornix (right panels) sampled at necropsy from PBS+DT (**B**) and IL-7+DT (**C**) immunized macaques were labeled with anti-CD3 (red), anti-CD20 (cyan) and anti-CD31 (yellow) antibodies. Nuclei were stained with DAPI (blue). *EP: Pluristratified Epithelium; Ch: Chorion.* **(D, E)** Sections (n=8 to 14 sections per macaque) of vaginal mucosa gathered from the PBS+DT (white boxes; Mac#1, #2 and #3) and the IL-7+DT (black boxes; Mac#4, #5 and #6) immunized macaques at necropsy were immunostained with anti-CD3 and anti-CD20 antibodies. The number of lymphoid follicles **(D)** and the percentage of B-cells in each follicle (n=7 to 21 follicles analyzed per macaque) **(E)** are presented as box-plots. Statistical differences between the 2 groups of macaques are shown (Mann-Whitney U test). **(F, G)** Sections of vaginal walls (left panels) and vaginal fornix (right panels) gathered from PBS+DT **(F)** and IL-7+DT **(G)** immunized macaques sampled at necropsy were labeled with anti-CD3 (red) and anti-CD20 (cyan) antibodies in combination with anti-PNAd (top panels), anti-GL-7 (middle panels) or anti-Ki-67 (bottom panels) (yellow) antibodies. Nuclei were stained with DAPI (blue). Arrows identify Ki-67-expressing B-cells.

The vaginal tissues taken at necropsy were analyzed by immunohistochemistry. In both groups of macaques, we demonstrated the presence of organized lymphoid follicles, composed of B- and T-cells located close to CD31^+^ endothelial cells (**Figures 7B-C**). However, in IL-7-treated DT-immunized macaques, these structures were both more numerous (11±2, 23±4, 16±2 follicles/50mm^2^ of tissue in IL-7-treated macaques and 8±2, 8±1, 6±1 follicles/50mm^2^ of tissue in control macaques; p<0.05; **Figure 7D**) and enriched in B lymphocytes (27±2%, 22±3%, 25±4% of B-cells in follicles of the IL-7-treated macaques and 20±5%, 11±4%, 16±3% of B-cells in follicles of the control macaques; p<0.05; **Figure 7E**), suggesting that their generation/maintenance was dependent on IL-7 stimulation.

In these follicles, PNAd^+^ (peripheral node addressin) high endothelial venule cells (**Figures 7F-G**, top panels) and GL-7^+^ T-cells were also in greater numbers (**Figures 7F-G**, middle panels). Interestingly, GL7^+^ B-cells were almost absent from the B cell zones, indicating follicles without organized germinal centers. However, while the vast majority of cycling (Ki-67^+^) cells were T-cells in macaques immunized with DT alone, both T- and B-cells were similarly cycling in the follicles of IL-7-treated DT-immunized macaques, suggesting ongoing local B-cell responses (**Figures 7F-G**, bottom panels, and **Supplementary Figure 5**, arrows indicate Ki-67^+^ B-cells).

Therefore, the pronounced increase in CCL19, CCL21, CXCL12 and CXCL13, together with the clustering of B- and T-cells in close proximity to endothelial cells expressing PNAd in the vaginal mucosa, indicates that pretreatment with rs-IL-7gly induces the formation of ectopic tertiary lymphoid follicles, which probably participate in the development of a stronger mucosal IgA immune response to DT.

## DISCUSSION

Similarly to what was observed in macaques subjected to systemic treatment with IL-7 (28), we demonstrated that local administration of rs-IL-7gly, either injected into or spayed onto the vaginal mucosa leads to local expression of a large array of chemokines within 48 hours following treatment. However, depending on the tissue responding to IL-7 (i.e. skin, intestine, lungs, vagina), the panel of overexpressed chemokines was different. In the IL-7-treated vaginal mucosa, 12 chemokines among 19 tested demonstrated increased expression either at the mRNA or at the protein levels, or both (**Figures 1** and **2**). Interestingly, the administered dose that was sufficient to drive chemokine expression in the vaginal mucosa was in the range of local IL-7 concentration observed in the ileum of acutely SIV-infected rhesus macaques (25) and after systemic injection of radiolabeled IL-7 to macaques (Cytheris S.A., now Revimmune Inc., personal communication).

Some of these chemokines (i.e. CCL2, CCL5, CCL17, CCL20, CXCL10 and CXCL12) are constitutively produced by cells of the FGT and participate in baseline immune cell turnover in the vaginal mucosa (5–8, 10). In contrast, local stimulation by CpG ODN or α-GalCer stimulates CCL2, CCL7, CCL19, CCL20, CCL22, CXCL8, CXCL10 or CX_3_CL1 overexpression in various mucosal models of inflammation, leading to the homing of immune cells into the mucosa (8, 37–39). Additionally, CCL28, which is expressed by diverse mucosal epithelia and selectively attracts IgA^+^ ASCs, is also driving the homing of antigen-specific cells into the vaginal mucosa (12).

However, one cannot exclude that some of these overexpressions of chemokine could also be indirectly stimulated by cytokines whose expression is triggered by IL-7 stimulation in the vaginal mucosa. Indeed, we evidenced an increased TSLP mRNA expression in vaginal biopsies collected after vaginal administration of 10 and 15μg of rs-IL-7gly (Figure S2), this cytokine being reported to stimulate CCL17 and CCL22 expression by CD11c^+^ mDCs (40).

Considering the wide range of chemokines that were overexpressed in the vaginal tissue, one can expect the migration of many immune cell types into this mucosa following IL-7 stimulation. Indeed, CD4^+^ and CD8^+^ T-cells, B-cells, NK-cells as well as CD11c^+^ mDCs and macrophages were clearly attracted to the vaginal chorion by day 2 following IL-7 administration. Interestingly, while lymphocytes were situated in the entire depth of the mucosa, most of the APCs, and in particular CD11c^+^DC-SIGN^+^ cells, were recruited just underneath the epithelium (**Figure 3A**). This particular localization could be attributed to CCL2-dependent recruitment as this chemokine is expressed by squamous vaginal epithelial cells and more specifically at the basolateral surface of primary endocervical epithelial cells (41) and, following stimulation with IL-7, was almost exclusively detected in the vaginal epithelium (**Figure 2**). Similarly, CCL7 and CCL5, which mostly recruit CCR2^+^ and CCR5^+^ cells, are also overexpressed in the vaginal epithelial layers of the FGT upon IL-7 stimulation (**Figure 2**), and may contribute to the peculiar localization of APCs in the IL-7-treated vaginal mucosa (**Figure 3**) (9).

In contrast, IL-7 dependent enhancement of CCL19, CXCL12 and CXCL13 was mainly observed in the vaginal chorion (**Figure 2**), suggesting their role in the recruitment of cells implicated in the adaptive immune response. Indeed, these chemokines allow the recruitment of CCR7^+^, CXCR4^+^ and CXCR5^+^ cells, including naïve B-cells and both CD4^+^ and CD8^+^ resting T-cells, which constitute the lymphoid infiltrate that characterized the IL-7-treated mucosa and TLS that we observed in the IL-7-treated immunized macaques (**Figure 7**).

To respond to IL-7, mucosal cells should express the specific receptor for this cytokine, a heterodimer protein composed of the IL-7Rα-chain (CD127) and the γc-chain (CD132). In addition to resting T-cells, various non-lymphoid cell types also express the IL-7 receptor (IL-7R). Indeed, in agreement with the literature that describes CD127 expression on epithelial and endothelial cells of diverse origins (34, 42–45), we identified, in the vaginal mucosa, CD127 expression on CD31^+^ endothelial cells (**Figure 2**) and, at a lower level, epithelial cells (**Figure 2D**, panels D1, D2 and D6 and Figure S3B-D). Interestingly, these cells produce significant levels of CCL2 and CXCL8 following *in vitro* IL-7 stimulation (46). Similarly, and in contrast with the classically observed down-regulation in T-cells, *in vitro* IL-7 stimulation was able to stimulate the up-regulation of CD127, by human aortic endothelial cells at the mRNA level (47). In this experimental model, IL-7 stimulation triggered the expression of CCL2 and cell adhesion molecules (ICAM-1 and VCAM-1) both at the mRNA level and at the protein level. In addition, an overexpression of CD132, the IL-7R beta chain, was also documented for endothelial cells of both blood and lymphatic vessels (48).

Finally, we demonstrated that both CD68^+^ pro-inflammatory “M1” macrophages and CD11c^+^CD163^+^ cells in the vaginal mucosa express CD127. The latter subset probably belongs to macrophages with a mixed “M1/M2” phenotype (Figure S3). As in humans, CD11c^+^CD11b^+^CD14^+^ FGT DCs lack CD163 expression (49) while CD1c^−^CD14^+^CD163^+^ FGT APCs expressing lower level of CD11c were classified as macrophages (50). In addition, in rhesus macaques, both CD68^+^ and CD163^+^ macrophages were identified in tissues from the FGT (39). Moreover, CD127 expression was previously reported for mouse intestinal macrophages (26), human CD68^+^ synovial macrophages (44) or human CD68^+^ and CD163^+^ macrophages in cardiac ventricular tissues sampled from patients with myocarditis (51), as well as *in vitro* monocyte-derived human macrophages (52). Similarly, vaginal CD11c^+^ dendritic cells also express CD127 (Figure S3), suggesting that they can participate in the mucosal response to IL-7 stimulation. In fact, IL-7 responsiveness of human monocytes, mDC and pDC was previously demonstrated by their capacity to produce CCL17, CCL22 and TSLP upon *in vitro* IL-7 stimulation (53–55). It is thus possible that DCs and macrophages, initially attracted in the mucosa, participate in the chemokine expression we observed in the IL-7 stimulated vagina and contribute to the immune cell homing into the vagina, in a positive feedback loop.

We then took advantage of the increased numbers of immune cells in the IL-7-treated vaginal mucosae to stimulate an antigen-specific immune response in this mucosa and clearly demonstrated the efficacy of rs-IL-7gly as an adjuvant to help the development of anti-DT mucosal antibody responses. In the animals vaccinated after local rs-IL-7gly stimulation, anti-DT mucosal antibody responses were indeed earlier, stronger and more persistent than in macaques immunized through administration of DT alone (**Figure 4**). More importantly, this mucosal immune response was largely composed of locally produced IgAs, as shown by the almost exclusive presence of DT-specific IgA plasma cells in the upper vagina and fornix of IL-7-treated DT-immunized macaques (**Figure 5**) and the lack of systemic IgA response in these macaques (**Figures 5** and **6**). In contrast, rs-IL-7gly stimulation prior to vaginal immunization allowed for the development of a systemic IgG response characterized by the presence of DT-specific IgG antibody secreting cells in the iliac LNs sampled at necropsy and in blood by the second week following primary immunization (**Figures 5** and **6**). However, DT-specific IgG ASCs were also detected in the iliac LNs of DT-alone immunized macaques, at least 2 weeks after the fourth immunization (i.e. at necropsy).

Interestingly, enhanced mucosal cellular immunity was demonstrated after topical administration of a modified IL-7 (IL-7 fused to the immunoglobulin Fc fragment - IL7-Fc) in systemically immunized mice (56). Surprisingly, in this study, native IL-7 was inefficient to trigger immune cell homing to the vagina. However, in the Choi *et al.* study, the administered IL-7 was non-glycosylated and administered by simply being deposited on the vaginal mucosa. It is possible that the velocity given by spray administration in our experiments allowed a better penetration of the cytokine across the mucus and the epithelial barrier in the IL-7-treated macaques, leading to improved efficacy. Moreover, we performed cervico-vaginal lavages before each spray, which could be important in reducing the amount of mucus at the epithelial surface and could also allow the cytokine to penetrate more easily into the mucosa.

Beside their classical homing function, chemokines such as CCL19, CCL21, CXCL12 and CXCL13 are also implicated, together with cytokines such as IL-17A (enhanced in IL-7-treated vaginal mucosa sampled 2 days following the administration of rs-IL-7gly, Figure S2) in the organization of TLS and germinal center formation (57, 58). At day 2 following rs-IL-7gly administration, most of the infiltrating immune cells were scattered in the chorion but lymphoid aggregates composed of T-cells, B-cells and APCs could also be observed in the vaginal mucosa (**Figure 3A**, bottom panels). However, at this time point, these aggregates, which did not contain clearly defined T- and B-cell zones, cannot be considered as organized lymphoid structures. In contrast, we observed such structures in the mucosa of IL-7-treated monkeys sampled at necropsy and were much less present in control macaques (**Figure 7D**). In both the upper part of the vaginal walls and the vaginal fornix, lymphoid follicles organized in distinct T-cell and B-cell areas containing proliferating cells were often surrounding CD31^+^ endothelial cells expressing PNAd, a marker that characterizes high endothelial venules, the portal of entry for T- and B-cells into TLS (**Figure 7**, (59)). However, at this step, we did not detect clear GL7^+^ B-cells in these structures while T-cells express this marker and proliferate, suggesting antigen-induced local activation.

Altogether, these data support the hypothesis that mucosal administration of rs-IL-7gly induces massive CXCR5^+^ cell recruitment at HEVs where PNAd and CXCL13 are expressed and initiates TLS neogenesis within vaginal tissue. High levels of IgAs in the vaginal secretions are produced by mucosally localized plasma cells as evidenced by reverse immunohistofluorescent staining. In the vagina of IL-7-treated macaques the mucosal overexpression of CXCL12 probably plays a role in the infiltration of DT-specific plasma cells (60, 61).

In this study, we showed that, in non-human primates, rs-IL-7gly sprayed in the vaginal lumen penetrates the mucosa and stimulates CD127^+^ intra-mucosal cells to produce a large array of chemokines that mobilize the mucosal immune system. IL-7 induced chemokine expression in the vaginal tissue triggers the recruitment of various immune cells, and the activation of mDCs, allowing for the generation of TLS underneath the vaginal epithelium and the development of a strong mucosal immune response following subsequent topical administration of antigen. These data suggest that non-traumatic administration of IL-7 could be used as a mucosal adjuvant to elicit vaginal antibody response and provide a very promising strategy to provide protection against sexually transmitted infections.

## CONFLICT OF INTEREST

The authors declare that the research was conducted in the absence of any commercial or financial relationships that could be construed as a potential conflict of interest.

## AUTHOR CONTRIBUTIONS

MR, SL, SFM, BCdM and AS performed the experiments. MR and RC designed the study and the experiments. ASDD and MB helped for the setting up of the ELISA. MR, SL, and RC analyzed and interpreted the data. MR and RC wrote the manuscript. SL, SFM, BCdM, AS, ASDD, MB, IBV, ACC, RC and MR discussed the results, commented the manuscript and approved the final version.

## FUNDING

This work was carried out in partial fulfillment of Sandrine Logerot’s PhD thesis at Université Paris Descartes, Paris, France S.L.’s PhD thesis was supported by a CIFRE (Convention Industrielle de Formation par la Recherche) fellowship co-funded by the French government and Cytheris S.A. (now Revimmune Inc.), and by Inserm-ANRS and Université Paris Descartes.

This work was supported by the ANRS (Agence Nationale de Recherches sur le SIDA et les Hépatites Virales), ANRT (Association Nationale de la Recherche et de la Technologie), Inserm, CNRS, Univeristé de Paris and Cytheris S.A. (now Revimmune Inc.). The funders had no role in study design, data collection and analysis, decision to publish, or preparation of the manuscript.

## ACKNOLEDGMENTS

The authors would like to thank Drs. Céline Gommet, Christophe Joubert and Nathalie Bosquet as well as the staffs of the Institut Pasteur (Paris, France) and Infectious Disease Models and Innovative Therapies (IDMIT, Fontenay-aux-Roses, France) Primate Centers. IDMIT was supported by French government “Programme d’Investissements d’Avenir” (PIA; ANR-11-INBS-0008). The authors greatly acknowledge Maryline Favier, Franck Letourneur and Pierre Bourdoncle, respectively heads of HistIM (histology and microdissection platform), GENOM’IC (genomic platform) and IMAG’IC (cell imagery platform) core facilities of the Institut Cochin. The authors thank Thomas Guilbert from IMAG’IC platform at the Institut Cochin for writing the routine for ImageJ image analysis software. The authors acknowledge Cytheris S.A. (now Revimmune Inc.), for providing the recombinant glycosylated simian IL-7 and Aptar pharma for providing the spray devices. We would like to thank Paul Belle, English-French Interpreter, for copyediting this article.

## Notes

### Competing Interest Statement

The authors have declared no competing interest.

## REFERENCES

1. Cuburu N, Kweon MN, Song JH, Hervouet C, Luci C, Sun JB, et al. Sublingual immunization induces broad-based systemic and mucosal immune responses in mice. Vaccine (2007) 25(51):8598–610. Epub 2007/11/13. doi: 10.1016/j.vaccine.2007.09.073. PubMed PMID: 17996991.

2. Czerkinsky C, Holmgren J. Topical immunization strategies. Mucosal immunology (2010) 3(6):545–55. Epub 2010/09/24. doi: 10.1038/mi.2010.55. PubMed PMID: 20861833.

3. Kozlowski PA, Cu-Uvin S, Neutra MR, Flanigan TP. Comparison of the oral, rectal, and vaginal immunization routes for induction of antibodies in rectal and genital tract secretions of women. Infection and immunity (1997) 65(4):1387–94. Epub 1997/04/01. PubMed PMID: 9119478; PubMed Central PMCID: PMC175144.

4. Kozlowski PA, Williams SB, Lynch RM, Flanigan TP, Patterson RR, Cu-Uvin S, et al. Differential induction of mucosal and systemic antibody responses in women after nasal, rectal, or vaginal immunization: influence of the menstrual cycle. Journal of immunology (2002) 169(1):566–74. Epub 2002/06/22. doi: 10.4049/jimmunol.169.1.566. PubMed PMID: 12077289.

5. Fichorova RN, Anderson DJ. Differential expression of immunobiological mediators by immortalized human cervical and vaginal epithelial cells. Biol Reprod (1999) 60(2):508–14. Epub 1999/01/23. PubMed PMID: 9916021.

6. Sharkey DJ, Macpherson AM, Tremellen KP, Robertson SA. Seminal plasma differentially regulates inflammatory cytokine gene expression in human cervical and vaginal epithelial cells. Mol Hum Reprod (2007) 13(7):491–501. Epub 2007/05/08. doi: gam028 [pii] 10.1093/molehr/gam028. PubMed PMID: 17483528.

7. Satthakarn S, Hladik F, Promsong A, Nittayananta W. Vaginal innate immune mediators are modulated by a water extract of Houttuynia cordata Thunb. BMC Complement Altern Med (2015) 15:183. Epub 2015/06/17. doi: 10.1186/s12906-015-0701-9 [pii]. PubMed PMID: 26077233; PubMed Central PMCID: PMC4466860.

8. Cremel M, Berlier W, Hamzeh H, Cognasse F, Lawrence P, Genin C, et al. Characterization of CCL20 secretion by human epithelial vaginal cells: involvement in Langerhans cell precursor attraction. Journal of leukocyte biology (2005) 78(1):158–66. PubMed PMID: 15831560.

9. Rancez M, Couedel-Courteille A, Cheynier R. Chemokines at mucosal barriers and their impact on HIV infection. Cytokine Growth Factor Rev (2012) 23(4-5):233–43. Epub 2012/06/26. doi: 10.1016/j.cytogfr.2012.05.010 S1359-6101(12)00037-8 [pii]. PubMed PMID: 22728258.

10. Wira CR, Rodriguez-Garcia M, Patel MV. The role of sex hormones in immune protection of the female reproductive tract. Nature reviews Immunology (2015) 15(4):217–30. Epub 2015/03/07. doi: 10.1038/nri3819. PubMed PMID: 25743222; PubMed Central PMCID: PMC4716657.

11. Zhou JZ, Way SS, Chen K. Immunology of Uterine and Vaginal Mucosae: (Trends in Immunology 39, 302-314, 2018). Trends in immunology (2018) 39(4):355. Epub 2018/03/14. doi: 10.1016/j.it.2018.02.006. PubMed PMID: 29530651; PubMed Central PMCID: PMC5880711.

12. Aldon Y, Kratochvil S, Shattock RJ, McKay PF. Chemokine-Adjuvanted Plasmid DNA Induces Homing of Antigen-Specific and Non-Antigen-Specific B and T Cells to the Intestinal and Genital Mucosae. Journal of immunology (2020) 204(4):903–13. Epub 2020/01/10. doi: 10.4049/jimmunol.1901184. PubMed PMID: 31915263; PubMed Central PMCID: PMC6994839.

13. Kelly KA, Chan AM, Butch A, Darville T. Two different homing pathways involving integrin beta7 and E-selectin significantly influence trafficking of CD4 cells to the genital tract following Chlamydia muridarum infection. American journal of reproductive immunology (2009) 61(6):438–45. Epub 2009/04/28. doi: 10.1111/j.1600-0897.2009.00704.x. PubMed PMID: 19392981; PubMed Central PMCID: PMC2888875.

14. Davila SJ, Olive AJ, Starnbach MN. Integrin alpha4beta1 is necessary for CD4+ T cell-mediated protection against genital Chlamydia trachomatis infection. Journal of immunology (2014) 192(9):4284–93. Epub 2014/03/25. doi: 10.4049/jimmunol.1303238. PubMed PMID: 24659687; PubMed Central PMCID: PMC3995848.

15. Johansson EL, Rudin A, Wassen L, Holmgren J. Distribution of lymphocytes and adhesion molecules in human cervix and vagina. Immunology (1999) 96(2):272–7. Epub 1999/05/08. doi: 10.1046/j.1365-2567.1999.00675.x. PubMed PMID: 10233705; PubMed Central PMCID: PMC2326729.

16. Parr MB, Parr EL. Interferon-gamma up-regulates intercellular adhesion molecule-1 and vascular cell adhesion molecule-1 and recruits lymphocytes into the vagina of immune mice challenged with herpes simplex virus-2. Immunology (2000) 99(4):540–5. Epub 2000/05/03. doi: 10.1046/j.1365-2567.2000.00980.x. PubMed PMID: 10792501; PubMed Central PMCID: PMC2327183.

17. Escario A, Gomez Barrio A, Simons Diez B, Escario JA. Immunohistochemical study of the vaginal inflammatory response in experimental trichomoniasis. Acta Trop (2010) 114(1):22–30. Epub 2009/12/23. doi: 10.1016/j.actatropica.2009.12.002. PubMed PMID: 20025844.

18. Bertley FM, Kozlowski PA, Wang SW, Chappelle J, Patel J, Sonuyi O, et al. Control of simian/human immunodeficiency virus viremia and disease progression after IL-2-augmented DNA-modified vaccinia virus Ankara nasal vaccination in nonhuman primates. Journal of immunology (2004) 172(6):3745–57. Epub 2004/03/09. doi: 10.4049/jimmunol.172.6.3745. PubMed PMID: 15004179.

19. Sui Y, Zhu Q, Gagnon S, Dzutsev A, Terabe M, Vaccari M, et al. Innate and adaptive immune correlates of vaccine and adjuvant-induced control of mucosal transmission of SIV in macaques. Proceedings of the National Academy of Sciences of the United States of America (2010) 107(21):9843–8. Epub 2010/05/12. doi: 10.1073/pnas.0911932107. PubMed PMID: 20457926; PubMed Central PMCID: PMC2906837.

20. Toka FN, Pack CD, Rouse BT. Molecular adjuvants for mucosal immunity. Immunological reviews (2004) 199:100–12. Epub 2004/07/06. doi: 10.1111/j.0105-2896.2004.0147.x. PubMed PMID: 15233729.

21. Hu K, Luo S, Tong L, Huang X, Jin W, Huang W, et al. CCL19 and CCL28 augment mucosal and systemic immune responses to HIV-1 gp140 by mobilizing responsive immunocytes into secondary lymph nodes and mucosal tissue. Journal of immunology (2013) 191(4):1935–47. Epub 2013/07/17. doi: 10.4049/jimmunol.1300120. PubMed PMID: 23858028.

22. Van Roey GA, Arias MA, Tregoning JS, Rowe G, Shattock RJ. Thymic stromal lymphopoietin (TSLP) acts as a potent mucosal adjuvant for HIV-1 gp140 vaccination in mice. European journal of immunology (2012) 42(2):353–63. Epub 2011/11/08. doi: 10.1002/eji.201141787. PubMed PMID: 22057556; PubMed Central PMCID: PMC3378695.

23. Shin H, Kumamoto Y, Gopinath S, Iwasaki A. CD301b+ dendritic cells stimulate tissue-resident memory CD8+ T cells to protect against genital HSV-2. Nat Commun (2016) 7:13346. Epub 2016/11/09. doi: 10.1038/ncomms13346. PubMed PMID: 27827367; PubMed Central PMCID: PMC5105190.

24. Lillard JW, Jr., Boyaka PN, Hedrick JA, Zlotnik A, McGhee JR. Lymphotactin acts as an innate mucosal adjuvant. Journal of immunology (1999) 162(4):1959–65. Epub 1999/02/11. PubMed PMID: 9973465.

25. Ponte R, Rancez M, Figueiredo-Morgado S, Dutrieux J, Fabre-Mersseman V, Charmeteau-de-Muylder B, et al. Acute Simian Immunodeficiency Virus Infection Triggers Early and Transient Interleukin-7 Production in the Gut, Leading to Enhanced Local Chemokine Expression and Intestinal Immune Cell Homing. Frontiers in immunology (2017) 8:588. Epub 2017/06/06. doi: 10.3389/fimmu.2017.00588. PubMed PMID: 28579989; PubMed Central PMCID: PMC5437214.

26. Zhang W, Du JY, Yu Q, Jin JO. Interleukin-7 produced by intestinal epithelial cells in response to Citrobacter rodentium infection plays a major role in innate immunity against this pathogen. Infect Immun (2015) 83(8):3213–23. Epub 2015/06/03. doi: 10.1128/IAI.00320-15 IAI.00320-15 [pii]. PubMed PMID: 26034215; PubMed Central PMCID: PMC4496619.

27. Sieling PA, Sakimura L, Uyemura K, Yamamura M, Oliveros J, Nickoloff BJ, et al. IL-7 in the cell-mediated immune response to a human pathogen. Journal of immunology (1995) 154(6):2775–83. Epub 1995/03/15. PubMed PMID: 7876548.

28. Beq S, Rozlan S, Gautier D, Parker R, Mersseman V, Schilte C, et al. Injection of glycosylated recombinant simian IL-7 provokes rapid and massive T-cell homing in rhesus macaques. Blood (2009) 114(4):816–25. PubMed PMID: 19351957.

29. Cimbro R, Vassena L, Arthos J, Cicala C, Kehrl JH, Park C, et al. IL-7 induces expression and activation of integrin alpha4beta7 promoting naive T-cell homing to the intestinal mucosa. Blood (2012) 120(13):2610–9. Epub 2012/08/17. doi: 10.1182/blood-2012-06-434779. PubMed PMID: 22896005; PubMed Central PMCID: PMC3460683.

30. Meier D, Bornmann C, Chappaz S, Schmutz S, Otten LA, Ceredig R, et al. Ectopic lymphoid-organ development occurs through interleukin 7-mediated enhanced survival of lymphoid-tissue-inducer cells. Immunity (2007) 26(5):643–54. Epub 2007/05/25. doi: 10.1016/j.immuni.2007.04.009. PubMed PMID: 17521585.

31. Timmer TC, Baltus B, Vondenhoff M, Huizinga TW, Tak PP, Verweij CL, et al. Inflammation and ectopic lymphoid structures in rheumatoid arthritis synovial tissues dissected by genomics technology: identification of the interleukin-7 signaling pathway in tissues with lymphoid neogenesis. Arthritis and rheumatism (2007) 56(8):2492–502. Epub 2007/08/01. doi: 10.1002/art.22748. PubMed PMID: 17665400.

32. Nayar S, Campos J, Chung MM, Navarro-Nunez L, Chachlani M, Steinthal N, et al. Bimodal Expansion of the Lymphatic Vessels Is Regulated by the Sequential Expression of IL-7 and Lymphotoxin alpha1beta2 in Newly Formed Tertiary Lymphoid Structures. Journal of immunology (2016) 197(5):1957–67. Epub 2016/07/31. doi: 10.4049/jimmunol.1500686. PubMed PMID: 27474071; PubMed Central PMCID: PMC4991245.

33. Ciccia F, Rizzo A, Maugeri R, Alessandro R, Croci S, Guggino G, et al. Ectopic expression of CXCL13, BAFF, APRIL and LT-beta is associated with artery tertiary lymphoid organs in giant cell arteritis. Annals of the rheumatic diseases (2017) 76(1):235–43. Epub 2016/04/22. doi: 10.1136/annrheumdis-2016-209217. PubMed PMID: 27098405.

34. Al-Rawi MA, Watkins G, Mansel RE, Jiang WG. The effects of interleukin-7 on the lymphangiogenic properties of human endothelial cells. Int J Oncol (2005) 27(3):721–30. Epub 2005/08/04. PubMed PMID: 16077922.

35. Sereti I, Dunham RM, Spritzler J, Aga E, Proschan MA, Medvik K, et al. IL-7 administration drives T cell-cycle entry and expansion in HIV-1 infection. Blood (2009) 113(25):6304–14. Epub 2009/04/22. doi: 10.1182/blood-2008-10-186601. PubMed PMID: 19380868; PubMed Central PMCID: PMC2710926.

36. Ribeiro Dos Santos P, Rancez M, Pretet JL, Michel-Salzat A, Messent V, Bogdanova A, et al. Rapid dissemination of SIV follows multisite entry after rectal inoculation. PLoS One (2011) 6(5):e19493. Epub 2011/05/17. doi: 10.1371/journal.pone.0019493 PONE-D-10-04131 [pii]. PubMed PMID: 21573012; PubMed Central PMCID: PMC3090405.

37. Lindqvist M, Navabi N, Jansson M, Samuelson E, Sjoling A, Orndal C, et al. Local cytokine and inflammatory responses to candidate vaginal adjuvants in mice. Vaccine (2009) 28(1):270–8. Epub 2009/10/06. doi: 10.1016/j.vaccine.2009.09.083 S0264-410X(09)01430-3 [pii]. PubMed PMID: 19800444.

38. Schenkel JM, Fraser KA, Vezys V, Masopust D. Sensing and alarm function of resident memory CD8(+) T cells. Nature immunology (2013) 14(5):509–13. Epub 2013/04/02. doi: 10.1038/ni.2568. PubMed PMID: 23542740; PubMed Central PMCID: PMC3631432.

39. Shang L, Duan L, Perkey KE, Wietgrefe S, Zupancic M, Smith AJ, et al. Epithelium-innate immune cell axis in mucosal responses to SIV. Mucosal immunology (2017) 10(2):508–19. Epub 2016/07/21. doi: 10.1038/mi.2016.62. PubMed PMID: 27435105; PubMed Central PMCID: PMC5250613.

40. Fontenot D, He H, Hanabuchi S, Nehete PN, Zhang M, Chang M, et al. TSLP production by epithelial cells exposed to immunodeficiency virus triggers DC-mediated mucosal infection of CD4+ T cells. Proc Natl Acad Sci U S A (2009) 106(39):16776–81. Epub 2009/10/07. doi: 10.1073/pnas.09073471060907347106 [pii]. PubMed PMID: 19805372; PubMed Central PMCID: PMC2757857.

41. Fahey JV, Schaefer TM, Channon JY, Wira CR. Secretion of cytokines and chemokines by polarized human epithelial cells from the female reproductive tract. Hum Reprod (2005) 20(6):1439–46. Epub 2005/03/01. doi: deh806 [pii] 10.1093/humrep/deh806. PubMed PMID: 15734755.

42. Reinecker HC, Podolsky DK. Human intestinal epithelial cells express functional cytokine receptors sharing the common gamma c chain of the interleukin 2 receptor. Proc Natl Acad Sci U S A (1995) 92(18):8353–7. Epub 1995/08/29. PubMed PMID: 7667294; PubMed Central PMCID: PMC41155.

43. Dus D, Krawczenko A, Zalecki P, Paprocka M, Wiedlocha A, Goupille C, et al. IL-7 receptor is present on human microvascular endothelial cells. Immunol Lett (2003) 86(2):163–8. Epub 2003/03/20. doi: S016524780300018X [pii]. PubMed PMID: 12644318.

44. Pickens SR, Chamberlain ND, Volin MV, Pope RM, Talarico NE, Mandelin AM, 2nd, et al. Characterization of interleukin-7 and interleukin-7 receptor in the pathogenesis of rheumatoid arthritis. Arthritis Rheum (2011) 63(10):2884–93. Epub 2011/06/08. doi: 10.1002/art.30493. PubMed PMID: 21647866; PubMed Central PMCID: PMC3614067.

45. Liao B, Cao PP, Zeng M, Zhen Z, Wang H, Zhang YN, et al. Interaction of thymic stromal lymphopoietin, IL-33, and their receptors in epithelial cells in eosinophilic chronic rhinosinusitis with nasal polyps. Allergy (2015) 70(9):1169–80. Epub 2015/06/23. doi: 10.1111/all.12667. PubMed PMID: 26095319.

46. Elner VM, Elner SG, Standiford TJ, Lukacs NW, Strieter RM, Kunkel SL. Interleukin-7 (IL-7) induces retinal pigment epithelial cell MCP-1 and IL-8. Exp Eye Res (1996) 63(3):297–303. Epub 1996/09/01. doi: S0014-4835(96)90118-9 [pii] 10.1006/exer.1996.0118. PubMed PMID: 8943702.

47. Li R, Paul A, Ko KW, Sheldon M, Rich BE, Terashima T, et al. Interleukin-7 induces recruitment of monocytes/macrophages to endothelium. Eur Heart J (2012) 33(24):3114–23. Epub 2011/08/02. doi: 10.1093/eurheartj/ehr245ehr245 [pii]. PubMed PMID: 21804111; PubMed Central PMCID: PMC3598429.

48. Iolyeva M, Aebischer D, Proulx ST, Willrodt AH, Ecoiffier T, Haner S, et al. Interleukin-7 is produced by afferent lymphatic vessels and supports lymphatic drainage. Blood (2013) 122(13):2271–81. Epub 2013/08/22. doi: 10.1182/blood-2013-01-478073 blood-2013-01-478073 [pii]. PubMed PMID: 23963040; PubMed Central PMCID: PMC3952712.

49. Rodriguez-Garcia M, Shen Z, Barr FD, Boesch AW, Ackerman ME, Kappes JC, et al. Dendritic cells from the human female reproductive tract rapidly capture and respond to HIV. Mucosal immunology (2017) 10(2):531–44. Epub 2016/09/01. doi: 10.1038/mi.2016.72. PubMed PMID: 27579858; PubMed Central PMCID: PMC5332537.

50. Duluc D, Gannevat J, Anguiano E, Zurawski S, Carley M, Boreham M, et al. Functional diversity of human vaginal APC subsets in directing T-cell responses. Mucosal immunology (2013) 6(3):626–38. Epub 2012/11/08. doi: 10.1038/mi.2012.104. PubMed PMID: 23131784; PubMed Central PMCID: PMC3568194.

51. Kubin N, Richter M, Sen-Hild B, Akinturk H, Schonburg M, Kubin T, et al. Macrophages represent the major pool of IL-7Ralpha expressing cells in patients with myocarditis. Cytokine (2020) 130:155053. Epub 2020/03/24. doi: 10.1016/j.cyto.2020.155053. PubMed PMID: 32203694.

52. Zhang M, Drenkow J, Lankford CS, Frucht DM, Rabin RL, Gingeras TR, et al. HIV regulation of the IL-7R: a viral mechanism for enhancing HIV-1 replication in human macrophages in vitro. Journal of leukocyte biology (2006) 79(6):1328–38. Epub 2006/04/15. doi: jlb.0704424 [pii] 10.1189/jlb.0704424. PubMed PMID: 16614257.

53. McKay FC, Hoe E, Parnell G, Gatt P, Schibeci SD, Stewart GJ, et al. IL7Ralpha expression and upregulation by IFNbeta in dendritic cell subsets is haplotype-dependent. PLoS One (2013) 8(10):e77508. Epub 2013/10/23. doi: 10.1371/journal.pone.0077508 PONE-D-12-30789 [pii]. PubMed PMID: 24147013; PubMed Central PMCID: PMC3797747.

54. Reche PA, Soumelis V, Gorman DM, Clifford T, Liu M, Travis M, et al. Human thymic stromal lymphopoietin preferentially stimulates myeloid cells. Journal of immunology (2001) 167(1):336–43. Epub 2001/06/22. doi: 10.4049/jimmunol.167.1.336. PubMed PMID: 11418668.

55. Vulcano M, Albanesi C, Stoppacciaro A, Bagnati R, D’Amico G, Struyf S, et al. Dendritic cells as a major source of macrophage-derived chemokine/CCL22 in vitro and in vivo. European journal of immunology (2001) 31(3):812–22. Epub 2001/03/10. doi: 10.1002/1521-4141(200103)31:3<812::AID-IMMU812>3.0.CO;2-L [pii] 10.1002/1521-4141(200103)31:3<812::AID-IMMU812>3.0.CO;2-L. PubMed PMID: 11241286.

56. Choi YW, Kang MC, Seo YB, Namkoong H, Park Y, Choi DH, et al. Intravaginal Administration of Fc-Fused IL7 Suppresses the Cervicovaginal Tumor by Recruiting HPV DNA Vaccine-Induced CD8 T Cells. Clin Cancer Res (2016) 22(23):5898–908. Epub 2016/07/14. doi: 10.1158/1078-0432.CCR-16-0423. PubMed PMID: 27407095.

57. Jones GW, Jones SA. Ectopic lymphoid follicles: inducible centres for generating antigen-specific immune responses within tissues. Immunology (2016) 147(2):141–51. Epub 2015/11/10. doi: 10.1111/imm.12554. PubMed PMID: 26551738; PubMed Central PMCID: PMC4717241.

58. Luo S, Zhu R, Yu T, Fan H, Hu Y, Mohanta SK, et al. Chronic Inflammation: A Common Promoter in Tertiary Lymphoid Organ Neogenesis. Frontiers in immunology (2019) 10:2938. Epub 2020/01/11. doi: 10.3389/fimmu.2019.02938. PubMed PMID: 31921189; PubMed Central PMCID: PMC6930186.

59. Ruddle NH. High Endothelial Venules and Lymphatic Vessels in Tertiary Lymphoid Organs: Characteristics, Functions, and Regulation. Frontiers in immunology (2016) 7:491. Epub 2016/11/25. doi: 10.3389/fimmu.2016.00491. PubMed PMID: 27881983; PubMed Central PMCID: PMC5101196.

60. Hargreaves DC, Hyman PL, Lu TT, Ngo VN, Bidgol A, Suzuki G, et al. A coordinated change in chemokine responsiveness guides plasma cell movements. The Journal of experimental medicine (2001) 194(1):45–56. Epub 2001/07/04. doi: 10.1084/jem.194.1.45. PubMed PMID: 11435471; PubMed Central PMCID: PMC2193440.

61. Hiepe F, Radbruch A. Plasma cells as an innovative target in autoimmune disease with renal manifestations. Nat Rev Nephrol (2016) 12(4):232–40. Epub 2016/03/01. doi: 10.1038/nrneph.2016.20. PubMed PMID: 26923204.

