## Supplemental materials for "IL-7-adjuvanted vaginal vaccine elicits strong mucosal immune responses in non-human primates"

*Supplementary Material*

### SUPPLEMENTARY MATERIALS AND METHODS

#### Immunohistofluorescent staining.

Four-µm thick tissue sections fixed in formaldehyde, and embedded in paraffin (FFPE), collected on glass slides (SuperFrost ® Plus, Menzel-Gläser, Illkirch, France) were dewaxed and rehydrated. Unmasking of the antigens was carried out by pressure cooking the sections for 15 minutes in 0.01M buffered sodium citrate solution (pH 6) or for 20 minutes in 1mM EDTA/10mM Tris Buffer (pH 9). Sections were then rinsed with PBS.

Eight-µm thick cryosections were collected on glass slides (SuperFrost® Plus), air dried overnight, fixed with 100% acetone for 10 minutes at room temperature (RT), air dried for 4 hours then stored at -80°C until use. Before immunostaining, frozen sections were fixed for 5 minutes at RT in 50/50 acetone/methanol and rinsed with PBS, excepted for chemokine staining.

Both types of sections were incubated for 30 minutes with blocking buffer (2% normal goat serum and 5% BSA in PBS or 5% BSA in PBS when using primary goat antibodies), incubated overnight with primary antibodies at 4°C, washed in PBS/0.5%Tween20 (Sigma-Aldrich) at RT, incubated with secondary antibodies in the dark for 30 minutes at RT and washed in PBS/0.5%Tween20. Finally, sections were washed in PBS alone, counterstained with 4,6-diamidino-2-phenylindol/DAPI (Molecular Probes, Cergy Pontoise, France) and mounted in Fluoromount-G (Southern Biotechnology, Birmingham, England). The antibodies used for immunohistofluorescence labeling are listed in Supplementary Table 2. For staining on FFPE sections, a supplemental 10 minutes incubation in 10mM $CuSO_4$/50mM $NH_4Cl$ (pH 5) at RT was added after the DAPI step to reduce autofluorescence. For CD127 staining on FFPE sections, a 30 minutes incubation step in 2% normal goat serum, 5% BSA, 10% FcR Blocking Reagent (Miltenyi Biotec, Paris, France) in PBS was added followed by an amplification step with a biotin-conjugated secondary antibody and a streptavidin conjugated to Alexa Fluor® 488, 546 or 633 (Molecular Probes) after having used a Streptavidin/Biotin Kit according to manufacturer's recommendations (Vector Laboratories, Nanterre, France). OPAL™ kit was used according to manufacturer's recommendations (PerkinElmer®, Courtaboeuf, France), to detect anti-CD127 in combination with CD3 and MHC-II MamuLa-DR. Of note, anti-CD127 was incubated before anti-CD3 and MHC-II MamuLa-DR, which were then detected with Alexa Fluor®-conjugated secondary antibodies (Molecular Probes).

#### Quantification of total and DT-specific IgGs and IgAs by enzyme-linked immunosorbent assay.

Total and DT-specific immunoglobulins (IgGs or IgAs) were quantified in CVLs using an in house ELISA. For total Ig quantifications, Nunc™ MaxiSorp™ ELISA plates were coated with dilutions of samples and several dilutions of Human IgA (Jackson ImmunoResearch, West Grove, USA; 400 to 0.8 ng/mL) or Monkey IgG (Rockland Immunochemicals, Gilbertsville, USA; 400 to 0.8 ng/mL) standards and incubated overnight at 4°C, washed in PBS/0.1%Tween20, blocked with 2% BSA in PBS/0.1%Tween20 for 2 hours at 37°C, washed in PBS/0.1%Tween20, then incubated with horseradish peroxidase (HRP) conjugated goat anti-monkey IgA (Abnova, 0.1ug/mL) or IgG antibodies (Rockland Immunochemicals;

0.07ug/mL) for 1 hour at 37°C. After several washes in PBS/0.1%Tween20, 3,3',5,5'-Tetramethylbenzidine/TMB peroxidase substrate solution was added (KPL, Maryland, USA) and incubated for 10 minutes at RT, then the reaction was stopped with 1M of $H_3PO_4$. Plates were read at 450nm (SpectraMaxTM384 PLUS ELISA Microplate Reader, Molecular Devices) and duplicate experiments were performed for each sample. Data were analyzed using SoftMax Pro software (5.0.1 version, Molecular Devices). Total IgA or IgG concentrations were determined by interpolation, using the calibration line of IgA or IgG standards, respectively. For quantification of specific Igs, ELISA plates were coated with DT (1µg/mL) and incubated overnight at 4°C, washed in PBS/0.1%Tween20, blocked with 2% BSA in PBS/0.1%Tween20 for 2 hours at 37°C, incubated with dilutions of each samples overnight at 4°C, washed in PBS/0.1%Tween20, then goat anti-monkey IgA- (Abnova) or IgG-HRP conjugated antibodies (Rockland Immunochemicals) were added for 1 hour at 37°C. Finally, after several washes in PBS/0.1%Tween20, peroxidase substrate was added (TMB from KPL) and incubated for 30 minutes at 37°C and the reaction was stopped with 1M of $H_3PO_4$. Plates were read and analyzed as mentioned above. Samples with a signal at least twice above the background were considered positive. Results are expressed as OD for specific anti-DT divided by the concentration of total IgA or IgG (mg/mL) in a given sample.

**Preparation of cells for ELISPOT assay.**

Iliac LN cells were isolated by crushing the organs through a 40 µm cell strainer to obtain a single cell suspension. The cells were then washed in RPMI 1640 medium containing 100 U/mL penicillin, 100 µg/mL streptomycin and 10% fetal calf serum (RPMI/PS/10% FCS), centrifuged, and cell numbers and viability were determined by trypan blue exclusion.

Cells were prepared from the lower FGT. Small pieces of tissues (<10mm$^3$) were incubated in RPMI/PS/Hepes 20mM supplemented with dispase II (2.6 U/ml, Sigma) 2h at 37°C, then washed in RPMI/PS/Hepes 20mM supplemented with 5mM EDTA, then in RPMI/PS/Hepes 20mM and each time the supernatant was removed through a 100µm cell strainer. The tissue pieces were then digested in RPMI/PS supplemented with collagenase VIII (2 mg/mL, Sigma) and DNase (20 U/mL, ROCHE) for 1h30 on a magnetic stirrer at 37°C, the supernatant decanted through a 100µm cell strainer was saved, and the collagenase/DNase treatment was repeated once with fresh medium. The cells were washed in RPMI/PS/10% FCS, passed through 70µm then 40µm cell strainers, centrifuged, and the numbers and viability of cells were determined by trypan blue exclusion.

**Quantification of antibody secreting cells by B-cells ELISPOT.**

Antibody secreting cells (ASCs) were assayed in Multiscreen HA plates (Merck Milipore, Molsheim, France) coated with DT (10µg/mL), overnight at 4°C. After washes with PBS, the reactive sites were blocked by incubation with RPMI/PS/10% FCS for 2 hours at 37°C. After washes, PBMCs or cell suspensions recovered from tissue were transferred into plates at $1.10^6$, $4.10^5$, $1.10^5$ cells per well in RPMI/PS/10% FCS and incubated for 40 hours at 37°C in 5% $CO_2$. After removal of the cells, the plates were washed again and the DT-specific antibodies were detected with either goat anti-monkey IgG-HRP (Rockland Immunochemicals) or goat anti-monkey IgA-HRP (Abnova) and incubated 4 hours at 37°C. Plates were washed and spots were detected by addition of AEC (ImmPACT AEC, VECTOR Laboratories). Spots were counted in an ELISPOT Reader (BIOREADER®-5000-F, BioSys GmbH, Serlabo, Entraigues, France). Spot numbers were reported as DT-specific ASCs per million of PBMCs.

### SUPPLEMENTARY FIGURES

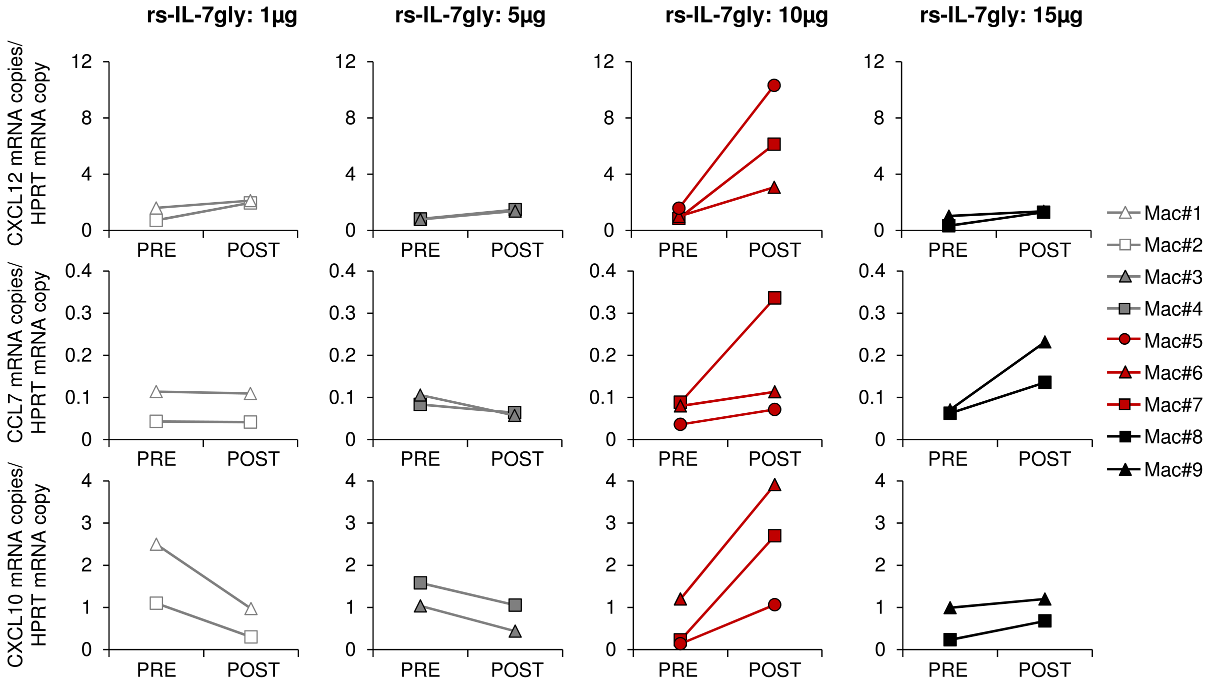


**Supplementary Figure 1. Topical rs-IL-7gly administration induces local chemokine transcription in the vaginal mucosa.**

The transcription of CXCL12 (top panels), CCL7 (middle panels) or CXCL10 (bottom panels) was quantified in vaginal biopsies sampled from macaques (n=9) one month before (PRE) and 48 hours after (POST) administration, by vaginal spray, of 1µg (light gray symbols), 5µg (dark gray symbols), 10µg (red symbols) or 15µg (black symbols) of rs-IL-7gly. Each point represents the median of 4 vaginal biopsies. For each sample, the data were normalized to HPRT mRNAs simultaneously quantified with the chemokines.

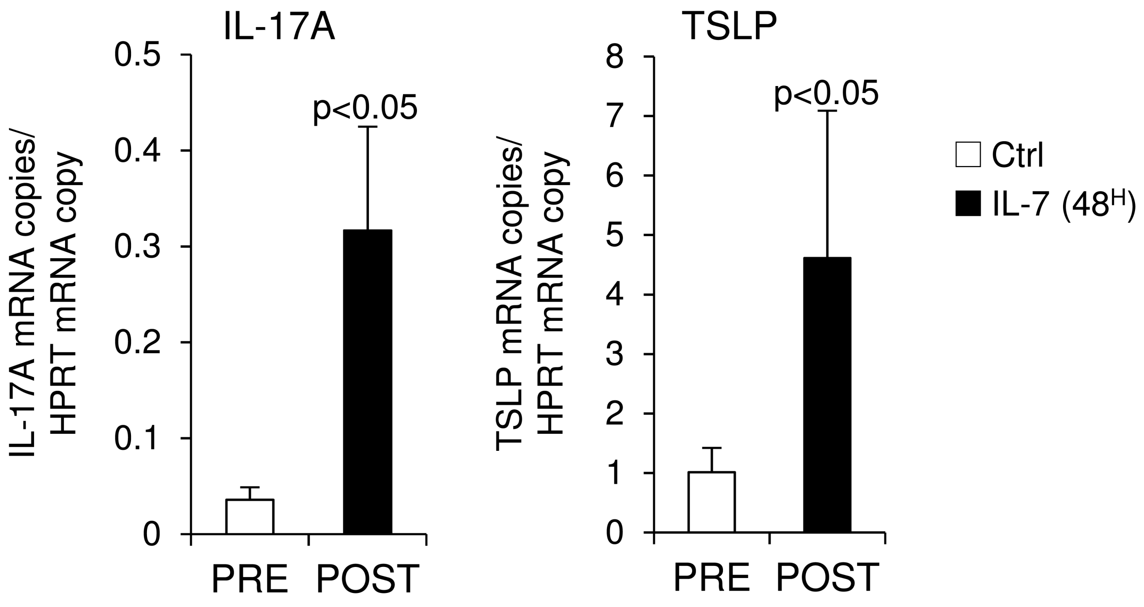


**Supplementary Figure 2. Topical administration of rs-IL-7gly induces local transcription of cytokines in the vaginal mucosa.**

The transcription of IL-17A and TSLP (Thymic stromal lymphopoietin) was quantified in vaginal biopsies (n=4 per macaque and time point) taken from macaques (n=5) one month before (Ctrl, white bars) and 48 hours after (IL-7 $48^H$, black bars) the administration of 10µg (n=3 macaques) or 15µg (n=2 macaques) of rs-IL-7gly, by vaginal spray, and normalized to HPRT mRNAs quantified simultaneously with the cytokines (mRNA copies/HPRT mRNA copy). Means and SEM of the data obtained for each of the 5 macaques are presented. Statistical differences are shown (Wilcoxon Signed-Rank Test).

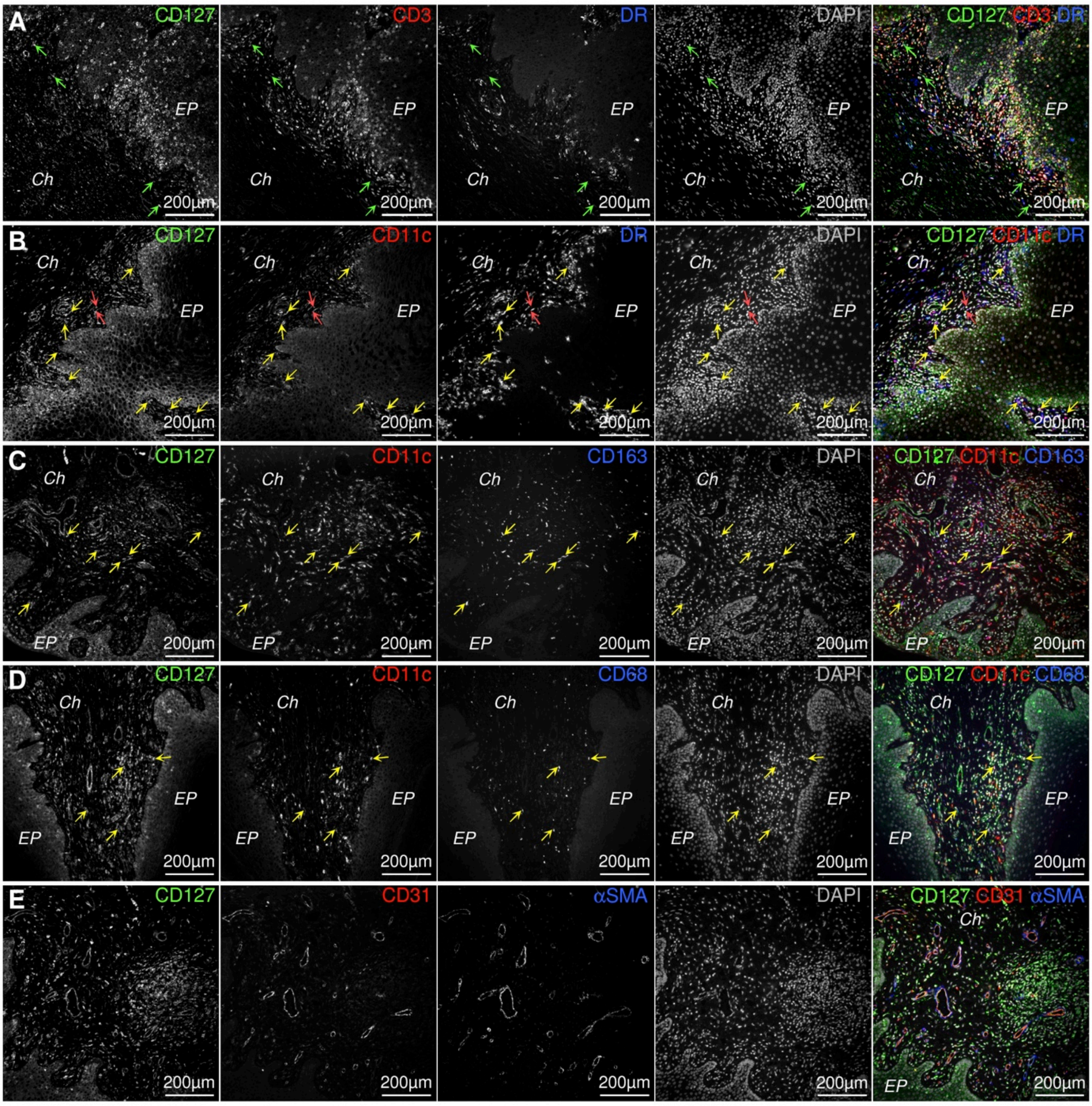


**Supplementary Figure 3. CD127-expressing cells in the vaginal mucosa.**

Sections of vaginal mucosa sampled from healthy monkeys were stained with anti-CD127 antibodies (**A-E**; green) in combination with anti-CD3 (**A**; red), anti-MamuLa-DR (**A, B**; blue), anti-CD11c (**B, C, D**; red), anti-CD163 (**C**; blue), anti-CD68 (**D**; blue), anti-CD31 (**E**; red) and/or anti-αSMA (**E**; blue) antibodies. Nuclei were stained with DAPI (grey). *Green arrows identify CD127$^+$CD3$^-$MamuLa-DR$^-$ cells; Yellow arrows identify triple positive cells: (B) CD127$^+$CD11c$^+$MamuLa-DR$^+$, or (C) CD127$^+$CD11c$^+$CD163$^+$, or (D) CD127$^+$CD11c$^+$CD68$^+$; Red arrows identify CD127$^+$CD11c$^+$MamuLa-DR$^-$ cells; EP: Pluristratified Epithelium; Ch: Chorion. DR: MHC-II MamuLa-DR.*

**A** – IL-7R and Vaginal Epithelial Cells

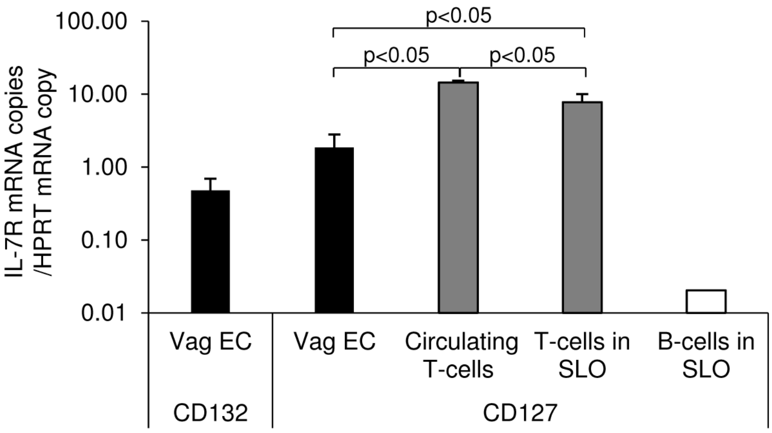


**B** – Adherent Vaginal Epithelial Cells in culture

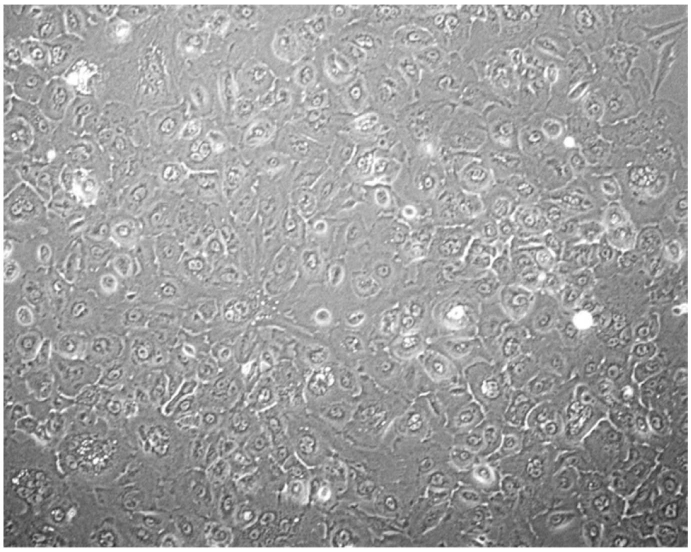

**Supplementary Figure 4. IL-7R and Vaginal Epithelial Cells.**

**(A)** mRNAs coding for CD132 (γc chain) and CD127 (IL-7Rα chain) were quantified in primary vaginal epithelial cells (Vag EC) obtained after Dispase/EDTA treatment of the vaginal epithelial layer, circulating T-cells, T-cells and B-cells purified from secondary lymphoid organ (SLO: spleen and lymph node) gathered from 3 healthy macaques and were normalized to HPRT mRNAs simultaneously quantified together with IL-7R mRNAs (IL-7R mRNA copies/HPRT mRNA copy). Bars and error bars represent means and SEM, respectively. Statistically significant differences are shown (Mann-Whitney U test). **(B)** Example of primary simian vaginal epithelial cells cultured for 2 weeks after isolation, on flask coated with collagen I in SmGM2 medium (Lonza).

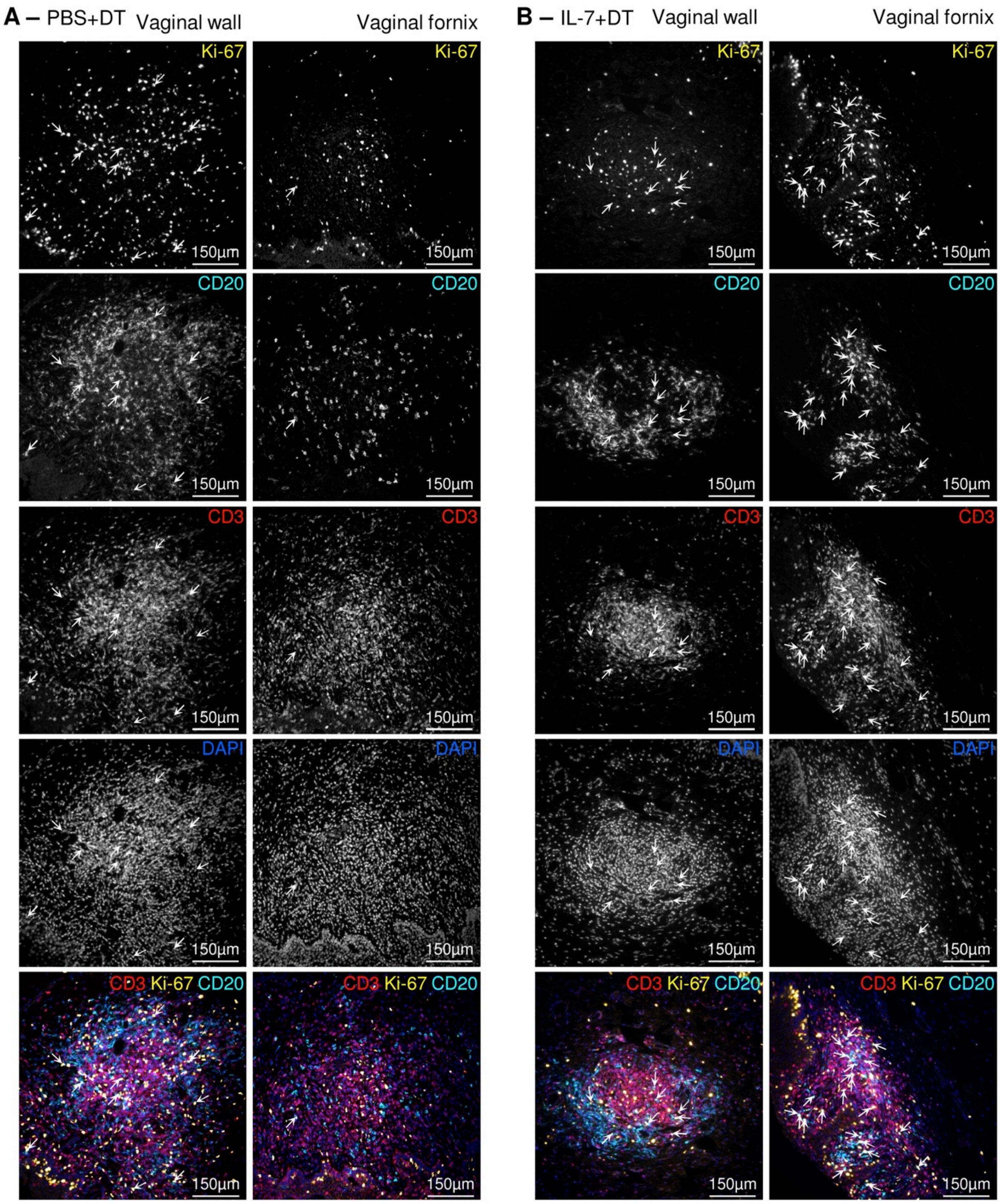


**Supplementary Figure 5. More B-cells proliferate in ectopic lymphoid follicles in the vaginal mucosa of IL-7-treated DT-immunized macaques.**

Sections of vaginal walls (left panels) and vaginal fornix (right panels) gathered from PBS+DT **(A)** and IL-7+DT **(B)** -immunized macaques at necropsy were labeled with anti-Ki-67 (yellow), anti-CD3 (red) and anti-CD20 (cyan) antibodies. Nuclei were stained with DAPI (blue). Arrows identify all B-cells expressing Ki-67.

### SUPPLEMENTARY TABLES

#### Supplementary Table 1. Oligonucleotides used for real-time PCR quantification of chemokines, cytokines and IL-7R mRNA.

| Outer 3'/5' primer pairs for first amplification | | Inner 3'/5' primer pairs for qPCR | |
|---|---|---|---|
| **Name** | **Sequence** | **Name** | **Sequence** |
| HPRT-Out5 | CTGAACGTCTTGCTCGAGAT | HPRT-In5 | CACATTGTAGCCCTCTGTGT |
| HPRT-Out3 | CGACCTTGACCATCTTTGGA | HPRT-In3 | CTGACCAAGGAAAGCAAAGT |
| CCL2-Out5 | AACATCCAGTGCTCAAACTGAA | CCL2-In5 | CTGCTCATAGCAGCCACCTTCA |
| CCL2-Out3 | TCCAGGTGGTCCATGGAAT | CCL2-In3 | TCCTGAACCCACTTCTGCTT |
| CCL3-Out5 | CATTCCATCACCTGCTCAGAA | CCL3-In5 | CAACCGGATCTCAGCAACAT |
| CCL3-Out3 | TCCAGGTCGCTGACGTATTT | CCL3-Int3 | GCCGGCCTCTCTTGGTTA |
| CCL4-Out5 | CCATGAAGCTGCGCGTGA | CCL4-In5 | CCCACCTCCTGCTGCTT |
| CCL4-Out3 | CTCAGTTCAGTTCCAGGTCA | CCL4-In3 | GCAGACTTGCTTCCCTCTTTT |
| CCL5-Out5 | GCTGTCATCCTCGTTGCTA | CCL5-In5 | ACATGCCTCAGACACCACA |
| CCL5-Out3 | CTCATCTCCAAAGAGTTGATGTA | CCL5-In3 | TACTCCCGAACCCATTTCTT |
| CCL7-Out5 | CATGCCCTCACCCTCCA | CCL7-In5 | CATGCCCTCACCCTCCA |
| CCL7-Out3 | CTCTGAGAAAGGACAGGGTA | CCL7-In3 | ATTTGTTTCTCAAGTCATGGCTT |
| CCL8-Out5 | GGACTTGCTCAGCCAGATT | CCL8-In5 | TGCTCAGCCAGATTCAGTTT |
| CCL8-Out3 | GGTCCAGATGCTTCATGGA | CCL8-In3 | CCTGACCCATCTCTCCTT |
| CCL11-Out5 | TGAAGGTCTCCACAACACT | CCL11-In5 | TGCAACCACCTGCTGCTTTA |
| CCL11-Out3 | TGGCTTTGGAGTTGGAGAT | CCL11-In3 | TTGGAGATTTTCGGTCCAGATA |
| CCL17-Out5 | GCAGCTCGAGGGACCA | CCL17-In5 | GCAGCTCGAGGGACCA |
| CCL17-Out3 | TCAAGGCTTTGCAGGTATTTAA | CCL17-In3 | GTTGGGGTCCGAACAGAT |
| CCL19-Out5 | ACCGTTGGCCTGCCTCTGTT | CCL19-In5 | CAGCCTGCTGGTTCTCTGGA |
| CCL19-Out3 | CTGCTGCGGCGCTTCATCTT | CCL19-In3 | TCCTCTGCAGTCTCTGGAT |
| CCL20-Out5 | CCATGTGCTGTACCAAGAGT | CCL20-In5 | TTGCTCCTGGCTGCTTTGA |
| CCL20-Out3 | TGCTGAGGCGACGTACAATA | CCL20-In3 | CCCAGGTCTGCTTTGGATT |
| CCL21-Out5 | CCTCAGCTCTGGCCTCTTA | CCL21-In5 | ATGGCTCAGTCACTGGCTCT |
| CCL21-Out3 | TCACTGGGCTATGGCCCTTTA | CCL21-In3 | CTATGGCCCTTTAGGGGTCT |
| CCL22-Out5 | CATGGCTCGCCTACAGA | CCL22-In5 | GTGACGCTTCAAGCAACTGA |
| CCL22-Out3 | GGGATGGAGATCAGGGAAT | CCL22-In3 | GATCAGGGAATGCAGAGAGT |
| CCL25-Out5 | CACACCCAAGGTGTCTTTGA | CCL25-In5 | TTATCGGATCCAGGAGGTGA |
| CCL25-Out3 | TTAGCTGATGTCAGGAGGGA | CCL25-In3 | CTGCTGGTGGGATTGCTAAA |
| CCL28-Out5 | AAGGCTCCTGGAAAGAGTGAA | CCL28-In5 | TCGCATCCAGAGAGCTGAT |
| CCL28-Out3 | GTTTCGTGTTTCCCCTGATGT | CCL28-In3 | CTGATGTGCCCTGTTACTGT |
| CXCL8-Out5 | GAACCATCTCGCTCTGTGTA | CXCL8-In5 | ATCTCGCTCTGTGTAAACATGA |
| CXCL8-Out3 | GCTCTCTTCCATCGGAAAGT | CXCL8-In3 | CTGTATTGGCACAGTGTGGT |
| CXCL10-Out5 | TGATTTGCTGCCTTGTCTTTCT | CXCL10-In5 | CTCTCTCAAGAACTGTACGCT |
| CXCL10-Out3 | ACCTCTTCTCACCCTTCTTTT | CXCL10-In3 | CATGTGGACAAAATTGACTTGGA |
| CXCL12-Out5 | GTCAGCCTGAGCTACAGAT | CXCL12-In5 | TACAGATGCCCATGCCGATT |
| CXCL12-Out3 | TTTCTCCAGGTACTCCTGAA | CXCL12-In3 | TCGGGTCAATGCACACTTGT |
| CXCL13-Out5 | TGCTTCTCATGCTGCTGGT | CXCL13-In5 | TCTCTCCAGTCCAAGGTGT |
| CXCL13-Out3 | AACACTGGAACTGGTGGAGT | CXCL13-In3 | GGAACTGGTGGAGTTGAAGA |
| CX3CL1-Out5 | TATCTCTGTCGTGGCTGCT | CX3CL1-In5 | GTCCTGCTGGCTGGACA |
| CX3CL1-Out3 | CCTCCATCCTGAGCCTTT | CX3CL1-In3 | TCCCTGAGGAACCAGTCA |
| IL-17A-Out5 | GCCATAGTGAAGGCAGGAAT | IL-17A-In5 | GCCATAGTGAAGGCAGGAAT |
| IL-17A-Out3 | AAGCCTAACTGCTTTGGGGA | IL-17A-In3 | AGTATCTTCTCCAGCCGGAA |

| | | | |
|---|---|---|---|
| IL-21-Out5 | CGTCTAGCTCTACTGTTGGT | IL-21-In5 | AGTCCTGGCAACATGGAGA |
| IL-21-Out3 | CAAGTTAGATCCTCAGGAATCTT | IL-21-In3 | CTAGAGGACAGATGCTGATGA |
| TSLP-Out5 | CTTTCAACTTGTAGGGCTGGT | TSLP-In5 | CTTTCAACTTGTAGGGCTGGT |
| TSLP-Out3 | TGTGACACTTGTTCCAGACATTT | TSLP-In3 | CTCTTCTTCATTGCCTGAGTA |
| LTb-Out5 | GATCAGGGAGGACTGGTAA | LTb-In5 | CAACAAGGACTGGGGTTTCA |
| LTb-Out3 | CGACGAGACAGTAGAGGTAA | LTb-In3 | CGACGAGACAGTAGAGGTAA |
| LTa-Out5 | CGTCAGCACCCCAAGAT | LTa-In5 | CAGAACTCACTGCTCTGGA |
| LTa-Out3 | ACAGTACTAGGGCTGAGGA | LTa-In3 | CCATCTGTGTGGGTGGATA |
| CD127-Out5 | TGCGCAAAATGGAGACTTGGA | CD127-In5 | GCCAGTTGGAAGTGAATGGA |
| CD127-Out3 | GTCATTGGCTCCTTCACGAT | CD127-In3 | TCAAAAGGAGCCTCAGGTTTAA |
| CD132-Out5 | CAGAAGTGCAGCCACTATCTA | CD132-In5 | CAGAAGTGCAGCCACTATCTA |
| CD132-Out3 | TTGTTCAGTCCAGCTGTGGT | CD132-In3 | GTGCTCCAAACAGTGGTTCA |

### Supplementary Table 2. Antibodies used for the immunostaining.

**Primary antibodies**

| **Target epitope, cell type, chemokine** | **Isotype** | **Clone** | **Supplier** | **Supplier details** |
|---|---|---|---|---|
| CD3 [a,b,c] | Rabbit IgG | (polyclonal) | DAKO | Trappes, France |
| CD4 [a] | Mouse IgG1 | L200 | BD Biosciences | Le Pont de Claix, France |
| CD8 [a] | Mouse IgG1 | RPA-T8 | BD Biosciences | Le Pont de Claix, France |
| CD20 [a,b,c] | Mouse IgG2a | L26 | DAKO | Trappes, France |
| CD31 [a,b,c$] | Mouse IgG1 | JC70A | DAKO | Trappes, France |
| PNAd [b,c] | Rat IgM | MECA-79 | Biolegend | St-Quentin en Yvelines, France |
| GL-7 [b,c] | Rat IgM | GL-7 | Biolegend | St-Quentin en Yvelines, France |
| HLA-DR [a,b] | Mouse IgG1 | TAL.1B5 | DAKO | Trappes, France |
| Tissue macrophages [a] | Mouse IgG1 | PM-2K | AbD Serotec | Düsseldorf, Germany |
| DC-SIGN [a] | Mouse IgG2b | 120507 | R&D systems | Lille, France |
| CD11c [a] | Mouse IgG1 | 3.9 | BioLegend | St-Quentin en Yvelines, France |
| CD11c [b] | Mouse IgG2a | 5D11 | LeicaBiosystems | Nanterre, France |
| CD68 [b] | Mouse IgG1 | KP1 | DAKO | Trappes, France |
| CD163 [b,] | Mouse IgG1 | 10D6 | Novacastra | Nanterre, France |
| αSMA [b,c$] | Mouse IgG2a | 1A4 | ThermoFischer | Villebon sur Yvette, France |
| CD83 [a] | Mouse IgG2b | HB15A | Beckman Coulter | Marseille, France |
| CD127 (C-ter) [b] | Rabbit | (polyclonal) | Abcam | Paris, France |
| Ki-67 [b] | Mouse IgG1 | B56 | BD Biosciences | Le Pont de Claix, France |
| CCL2 [a] | Rabbit | (polyclonal) | Aviva Systems Biology | San Diego, USA |
| CCL5 [a] | Rabbit | (polyclonal) | Aviva Systems Biology | San Diego, USA |
| CCL7 [a] | Rabbit | (polyclonal) | Aviva Systems Biology | San Diego, USA |
| CCL19 [a] | Mouse IgG2b | 54909 | R&D systems | Lille, France |
| CXCL13 [a] | Goat | (polyclonal) | R&D systems | Lille, France |
| CXCL12 [a] | Mouse IgG1 | 79018 | R&D systems | Lille, France |
| IgG [a] | Rabbit | (polyclonal) | DAKO | Trappes, France |
| IgA [a] | Rabbit | (polyclonal) | DAKO | Trappes, France |
| Anti-DT-FITC [a] | Goat IgG | (polyclonal) | Abcam | Paris, France |

**Secondary antibodies**

| **Target species** | **Host species** | **Conjugate** | **Supplier** | **Supplier details** |
|---|---|---|---|---|
| Mouse (IgG1, 2a or 2b) | Goat | Alexa Fluor® 546, 488 or 633 | Molecular Probes | Cergy Pontoise, France |
| Rabbit | Goat | Alexa Fluor® 546, 488 or 633 | Molecular Probes | Cergy Pontoise, France |
| Goat | Chicken | Alexa Fluor® 488 | Molecular Probes | Cergy Pontoise, France |
| Rabbit | Donkey | Alexa Fluor® 546 | Molecular Probes | Cergy Pontoise, France |
| Rabbit | Goat | Biotin | DAKO | Trappes, France |
| Goat | Donkey | Biotin | Abcam | Paris, France |

The tissue sections used for immunohistofluorescence labeling were [a] cryopreserved, [b] formaldehyde-fixed paraffin-embedded (FFPE) with acidic antigen retrieval (0.01M sodium citrate buffer, pH 6.0) or [d] FFPE with basic antigen retrieval (1mM EDTA/10mM Tris Buffer, pH 9.0). [$] Cell staining was much better and reproducible with basic antigen retrieval as compare as acidic antigen retrieval.
